# A Mathematical Model Connecting Appetite-Driven Food Intake to Metabolic Changes across Diet and GLP-1RA Treatments in Mice

**DOI:** 10.64898/2026.11.12.766168

**Authors:** Oscar Silfvergren, Christian Simonsson, Monika Sundqvist, Pär Nordell, Peter Gennemark, Gunnar Cedersund

## Abstract

Mouse models are widely used to study obesity and the effects of glucagon-like peptide-1 receptor agonists (GLP-1RAs) on appetite-driven meal responses and eating behaviours. However, connecting these behavioural changes to disease prevention remains limited. One reason is that eating behaviour is often summarised as daily or weekly energy intake, obscuring meal-level dynamics. Here we present a mathematical model of mouse appetite-driven food intake across diets and treatment with the GLP-1RAs semaglutide and exenatide. The model jointly described three studies and passed both qualitative assessments and a statistical χ² test (disagreement<threshold, 348<1111). Predictive performance was subsequently validated using four pre-existing studies and one new study. Finally, the model was used to simulate mouse populations under dietary and GLP-1RA interventions, with predictions showing qualitative agreement with the literature. This work aids preclinical drug discovery, supports mouse study design, and improves understanding of how individual meals accumulate into long-term physiological changes.

**Graphical abstract:** 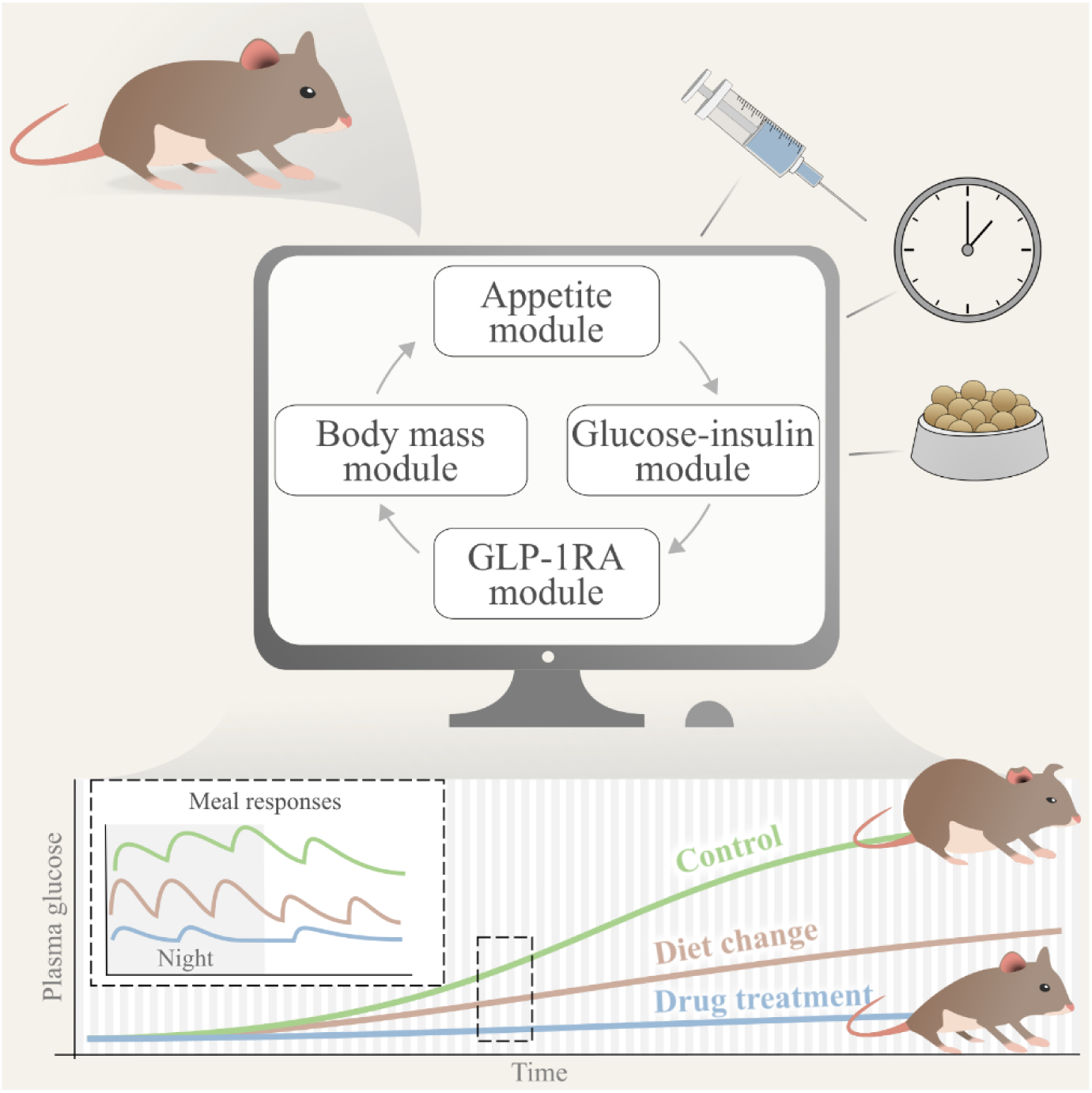

## INTRODUCTION

Rodent models, particularly mice, have been the gold standard for drug discovery and disease research for several decades (1,2). Since the early 1990s, following the discovery and development of the glucagon-like peptide-1 receptor agonist (GLP-1RA) exenatide, mice have been extensively studied to investigate the relationship between appetite inhibition and prevention of obesity-associated diseases such as diabetes. A more recent development is the GLP-1RA semaglutide, which has been reported to reduce body mass in diet-induced obese (DIO) mice up to 22% over a 3-week treatment period (3). Importantly, in these studies, mice are often given *ad libitum* access to food, providing essential insights into eating behaviours that reflect appetite, such as meal size, meal timing, and eating frequency. However, despite their use as the gold standard for several decades, our ability to connect dietary and GLP-1RA effects to these eating behaviours, and to downstream effects on body weight and disease prevention, remains limited. There are several reasons for this: connecting eating behaviours across mouse studies is challenging due to substantial heterogeneity in mouse strains and dietary conditions (4,5); GLP-1RAs have multi-timescale mechanisms of action (e.g., rapid increases in stimulated insulin secretion and slower accumulation of changes in body mass) that are difficult to describe mechanistically (6–9); and appetite-driven individual meals have historically often been summarised as daily or weekly energy averages, resulting in a loss of information (10,11). Addressing these challenges could improve mouse study design, aid our understanding of how individual meals accumulate into obesity-associated diseases, and guide the community towards more informed preclinical drug discovery. A powerful tool that could potentially address all these challenges is computational modelling.

A computational model is a mathematical abstraction of a system (12–14), and such models have been used for many years in drug discovery (15–17) and disease research (18–20). One type of computational model is an ordinary differential equation (ODE) model, which has been used to connect daily calorie intake to body mass changes in mice (21) and associated pharmaceutical effects (22). In 2023, that model was further developed by integrating a glucose-insulin model (23), resulting in a model connecting glucose-related metabolic dysregulation to diet-induced changes in body mass (24). While these models describe changes in body mass caused by changes in energy intake and expenditure, they do not describe eating behaviours or pharmaceutical effects on glucose-insulin regulation, such as those induced by GLP-1RAs. More recently, we developed a pharmacokinetic and pharmacodynamic (PKPD) model describing the effects of the GLP-1RA exenatide on glucose-insulin regulation (25). In addition to its effects on stimulated insulin secretion, the model accounted for pharmaceutical effects on appetite by decreasing meal sizes within a repeating meal schedule consisting of identical meals, timings, and eating frequencies. Taken together, these models provide important components for linking diet and GLP-1RA effects to body mass changes and disease prevention. One key component is still missing: a model of appetite-driven food intake that can describe feeding behaviours under *ad libitum* access to food and provide meals as inputs to the other mathematical models.

We present, to the best of our knowledge, a first mathematical model of mouse eating behaviour that incorporates cumulative dietary and GLP-1RA effects across meal responses and metabolic changes (**Fig. 1**). The model consists of a new appetite model connected to the pre-existing body mass (21,22), glucose-insulin (24), and GLP-1RA (25) models (**Fig. 1A**). Each of these models represents a particular subsystem of metabolism. The models can be simulated individually or in tandem. Therefore, these four models can be referred to as modules, which together constitute the full model (26). The appetite module describes meal consumption, the body mass module describes changes in body mass and fat mass, the GLP-1RA module describes the pharmacokinetics and downstream effects of the administered compound, and the glucose-insulin module describes glucose-insulin regulation. The resulting model’s ability to describe training data and predict independent studies (**Fig. 1B**) was supported by qualitative evaluations and statistical χ² tests across seven pre-existing studies. To further evaluate the model, we conducted a new semaglutide study, for which the model successfully predicted changes in appetite during and following GLP-1RA treatment. Finally, the model was used to generate digital mouse populations under various dietary and GLP-1RA treatment interventions, with predictions showing qualitative agreement with the literature (**Fig. 1B**). This work supports mouse study design in accordance with the principles of replacement, reduction, and refinement of animal use (27), improves our understanding of how individual meals accumulate to drive long-term changes, and guides the community towards more informed preclinical drug discovery.

**Figure 1.**
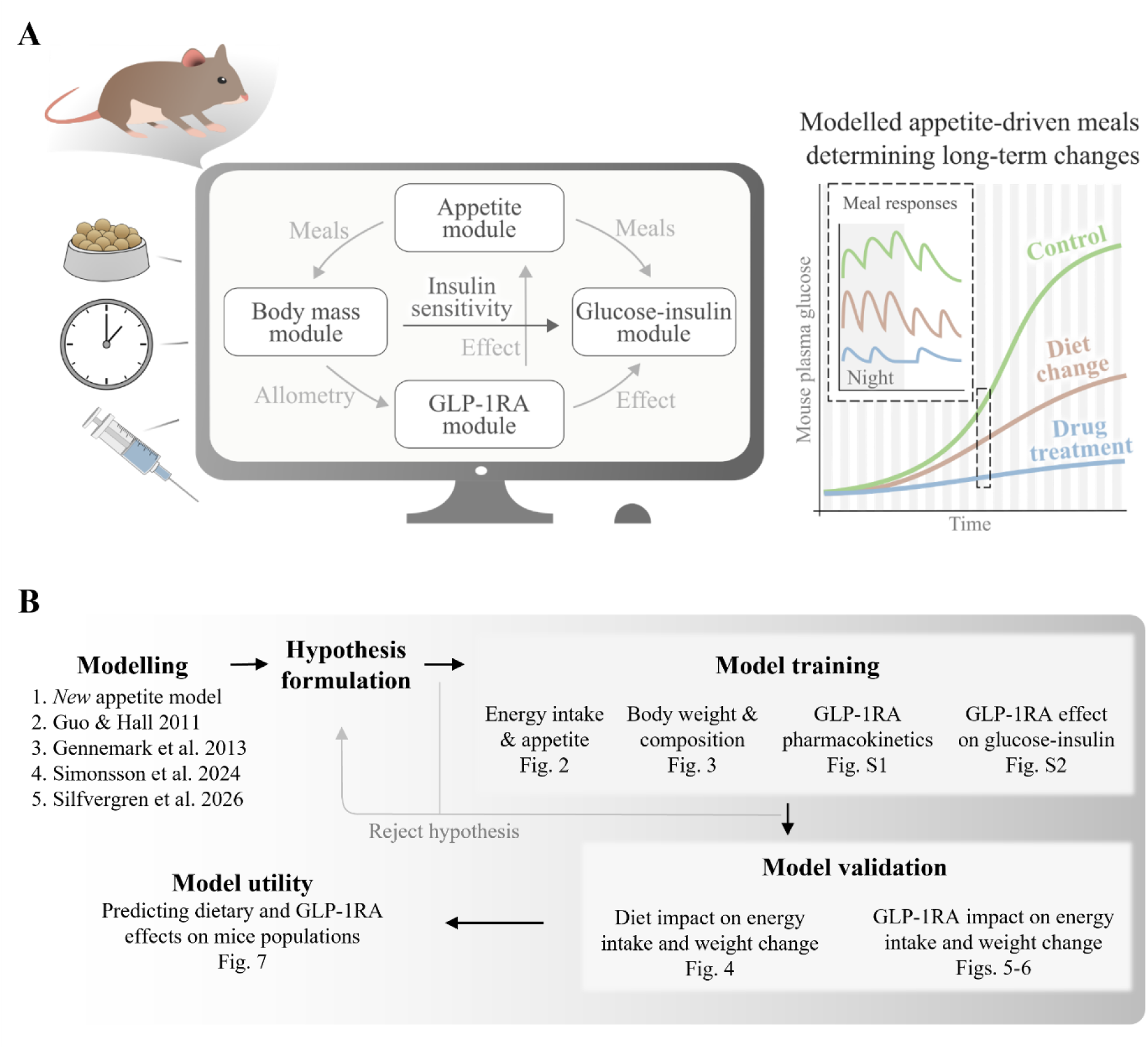
Integrated model architecture and study workflow. **(A)** Modular mathematical framework linking a novel appetite model with GLP-1RA treatment, glucose-insulin regulation (24,25), and body mass dynamics (21,22) in mice under ad libitum food access. **(B)** Overview of the study workflow, including modelling, hypothesis formulation, model training, model validation, and model utility.

## RESULTS

The model was evaluated in three stages. First, the model was trained to describe appetite-driven food intake and the resulting changes in body mass and metabolic regulation. Second, the model was validated using independent data on dietary interventions and treatment with the GLP-1RAs exenatide and semaglutide. Finally, the model was used to predict the effects of dietary and GLP-1RA interventions in healthy and obese mice. The model showed sufficient agreement with the training data based on qualitative evaluations and a statistical χ² test (joint sum weighted least squares with a minimum error 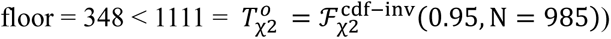

### Model training: Appetite module development and agreement with energy intake data

The appetite module describes energy intake across different diets and interventions (**Fig. 2**). It regulates food intake using a simplified description of the digestive system and a dynamic phenomenological variable termed *appetite* (**Fig. 2A**). Appetite is regulated by four key factors: i) circadian rhythm, with faster appetite development during the night and a gradual increase in amplitude over the course of the study to account for adaptation to the experimental setting and the associated increase in body mass; ii) fed state, through which mice gradually regulate food intake according to its caloric content; iii) dietary macronutrient composition, with lipids and non-lipids differing in their effects on satiety; and iv) GLP-1RA plasma exposure, which suppresses appetite according to a sigmoidal exposure–response relationship. When appetite exceeds a phenomenological threshold termed *hunger threshold*, meal consumption is initiated and continues until appetite falls below this threshold. Digested carbohydrates are transferred to the glucose-insulin module, where they appear in plasma, while the total energy leaving the digestive system is transferred to the body mass module as energy intake. The module comprises six ODEs and eight optimised parameters.

**Figure 2.**
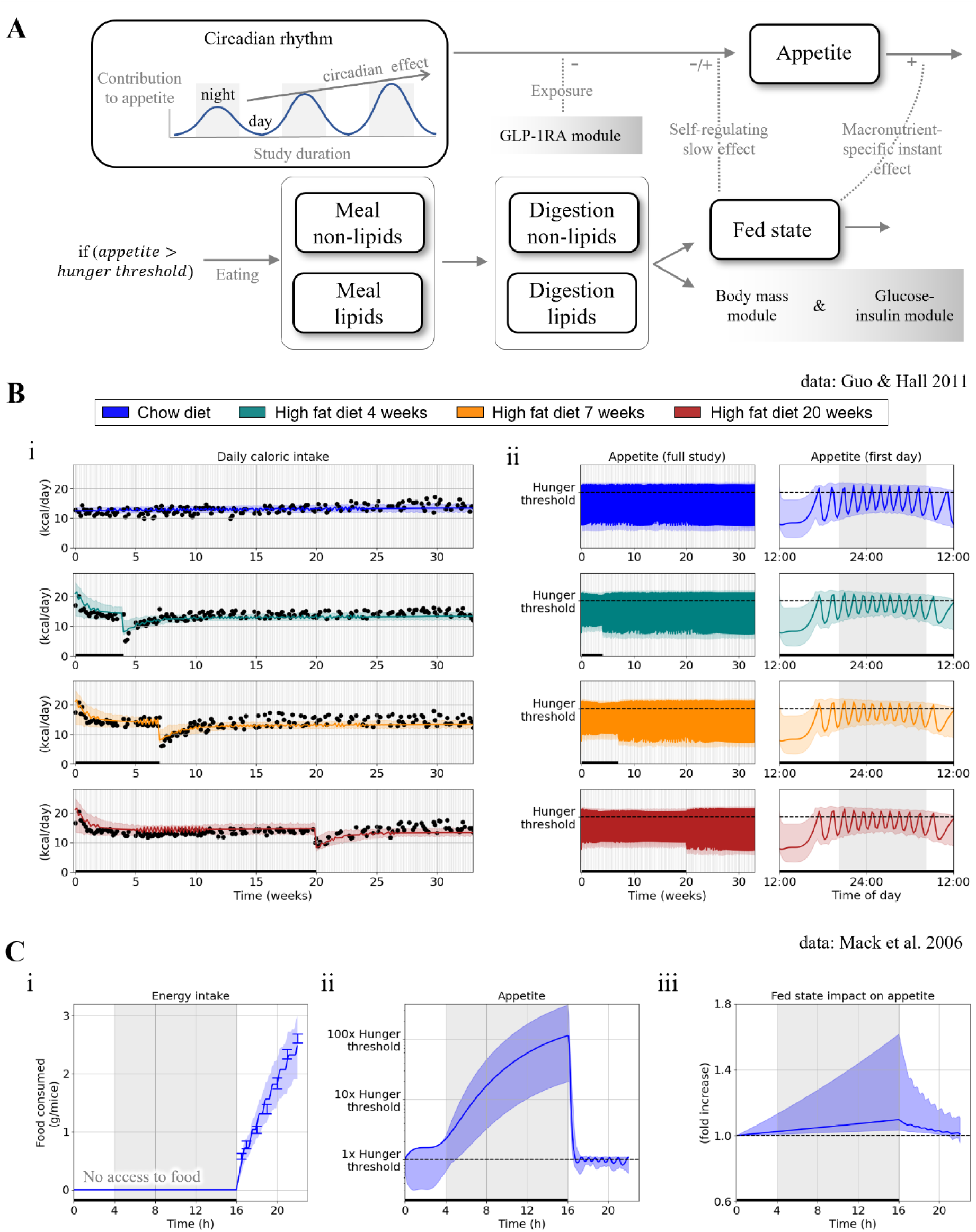
Model agreement with energy intake data. Lines represent the best fit of the model to all estimation data jointly (Figs. 2–3 and **Figs. S1–S2**), coloured areas indicate model uncertainty, grey areas indicate night-time, and black dots represent data (N = 1). **(A)** Illustration of the appetite module and its crosstalk with the other three modules. **(B)** Model description of a mouse study (21) in which mice were maintained on chow (blue) or switched from chow to a high-fat diet (HFD) and subsequently returned to chow after 4 weeks (turquoise), 7 weeks (orange), or 20 weeks (red). Black lines on the x-axis indicate periods of HFD feeding. **(i)** Energy intake data and model agreement with all estimation data jointly. **(ii)** Predicted appetite and hunger threshold for simulated meal consumption. **(C)** Model description of a mouse study (28) in which mice were deprived of food for 16 hours before receiving a standard chow diet (Teklad LM-485). The black line on the x-axis indicates the period without access to food. **(i)** Food consumption and model agreement with all estimation data jointly. **(ii)** Predicted appetite. **(iii)** Predicted fed-state regulation of appetite.

The model’s ability to describe energy intake was evaluated in mice maintained on chow diet (**Fig. 2B**, blue), or switched from chow to high-fat diet and subsequently back to chow at different time points (**Fig. 2B**, green, orange, and red) (21). Both simulations and experimental data showed a rapid increase followed by a slower decline in daily energy intake after switching to a high-fat diet, with the opposite pattern after returning to chow (**Fig. 2B i**, black bar). The model was further used to predict group-specific changes in appetite over the full study (**Fig. 2B ii**, left) and during the first 24 h of the study (**Fig. 2B ii**, right), where meal initiation is observed when *appetite* exceeds the marked *hunger threshold*.

The model’s ability to describe food intake following fasting was evaluated in mice fasted for 16 h followed by 6 h of *ad libitum* standard chow diet (Teklad LM-485) access (**Fig. 2C**) (28). Both simulations and experimental data showed frequent, high food intake during the refeeding period, with seven predicted meals and the largest meal immediately after fasting (**Fig. 2C i**). Model prediction uncertainty indicated that all mice would have initiated feeding approximately 5 h into the fast if food had been available (**Fig. 2C ii**). Appetite was predicted to increase throughout fasting, reaching 1.1-fold baseline (uncertainty range: 1.05–1.6), before returning toward baseline during refeeding (**Fig. 2C iii**).

### Model training: Agreement with body mass and metabolic dysregulation data

We next evaluated the model’s ability to describe long-term changes from individual meals (**Fig. 3**). The body mass module (**Fig. 3A**) used parameter values from the latest model version considered herein (24), except for physical activity-related energy expenditure, which was re-estimated to account for between-study variability. Although alternative parameterisations are possible, estimating physical activity has precedent in the literature and provided a practical way to capture between-study differences (21,22).

**Figure 3.**
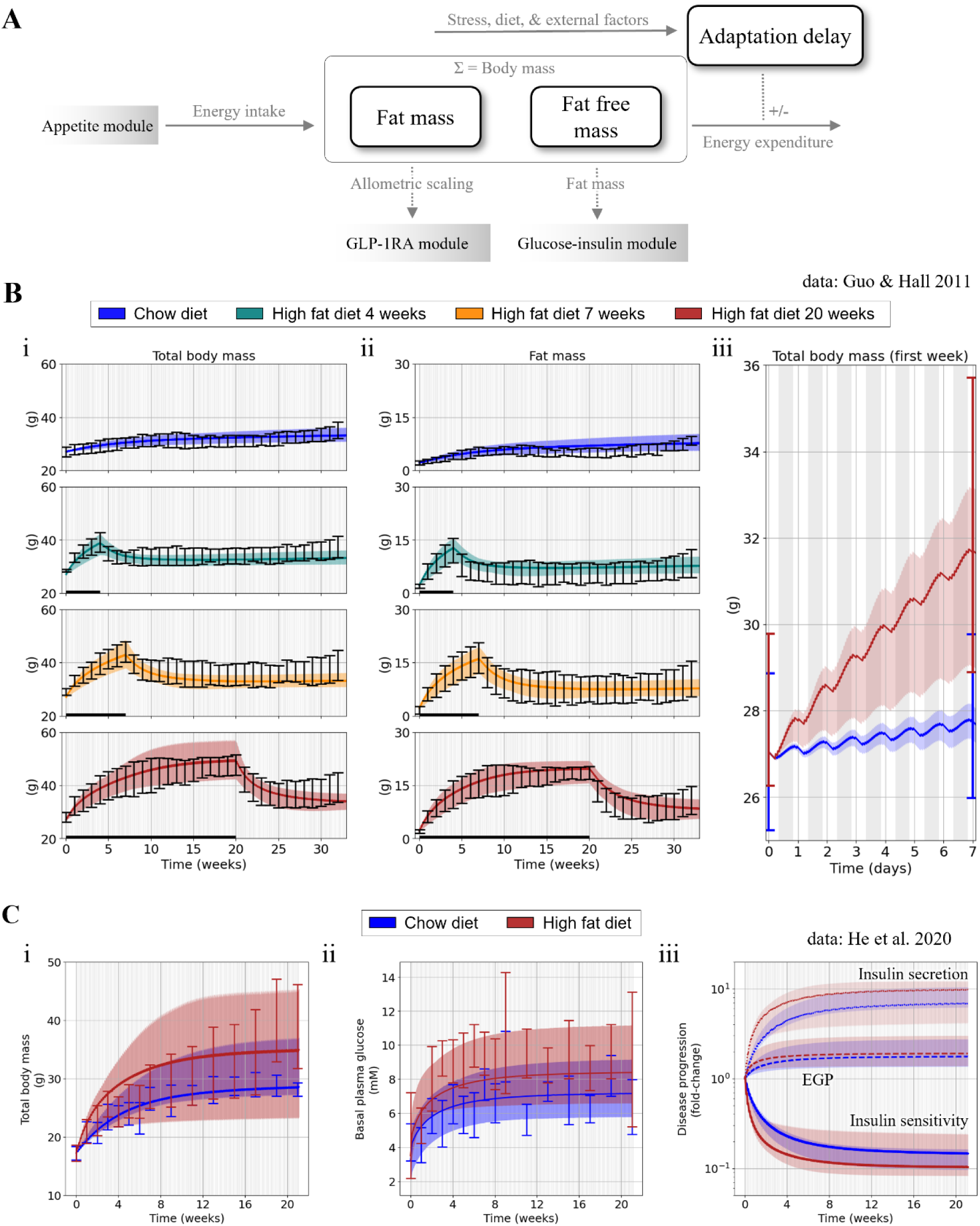
Model agreement with body mass changes and basal glucose levels. Lines represent the best fit of the model to all estimation data jointly (Figs. 2–3 and **Figs. S1–S2**), coloured areas indicate model uncertainty, grey areas indicate night-time, and error bars show population data (mean ± SEM). **(A)** Illustration of the body mass module and its crosstalk with the other three modules. **(B)** Model description of a mouse study (21) in which mice were maintained on chow (blue) or switched from chow to a high-fat diet (HFD) and subsequently returned to chow after 4 weeks (turquoise), 7 weeks (orange), or 20 weeks (red). Black lines on the x-axis indicate periods of HFD feeding. **(i)** Total body mass data and model agreement with all estimation data jointly. **(ii)** Fat mass data and model agreement with all estimation data jointly. **(iii)** Zoomed-in view of the study’s first week. **(C)** Model description of a mouse study (29) in which mice received either a chow diet (D12450J; blue) or a high-fat diet (D12492; red). **(i)** Total body mass data and model agreement with all estimation data jointly. **(ii)** Basal plasma glucose data and model agreement with all estimation data jointly. **(iii)** Predicted metabolic changes described by three variables: insulin secretion (dots), endogenous glucose production (EGP; dotted lines), and insulin sensitivity (solid lines).

The model’s ability to describe body mass changes was evaluated using experimental data from the same study in which the original model was presented (21) (**Fig. 3B**). The corresponding energy-intake data were also used for parameter estimation (**Fig. 2B**). Both simulations and experimental data showed rapid initial body mass gain followed by slower increases on a high-fat diet, with the opposite pattern after switching to chow (**Fig. 3B i**). Fat mass followed a similar pattern (**Fig. 3B ii**). Importantly, the model describes body mass accumulation as a cumulative consequence of meal intake, with greater increases at night when feeding is more frequent (**Fig. 3B ii**, grey area).

The model’s ability to describe cumulative effects of repeated meals was further evaluated using a study in which mice received *ad libitum* access to either chow (D12450J) or high-fat diet (D12492) (**Fig. 3C**) (29). Both experimental data and simulations showed greater body mass increases in mice fed the high-fat diet (**Fig. 3C i**). A similar pattern was observed in basal plasma glucose (**Fig. 3C ii**), which is an established biomarker of obesity-induced metabolic dysregulation (30,31). The model describes this dysregulation using three model variables implemented from the original glucose-insulin model (24): increased body fat increases insulin secretion (**Fig. 3C iii**, dots), increases endogenous glucose production (EGP) (**Fig. 3C iii**, dotted lines), and decreases insulin sensitivity (**Fig. 3C iii**, solid lines).

### Model validation: Predicting body mass changes and energy intake during repeated diet changes

The model’s ability to predict independent data was evaluated using a mouse group from the previously described Guo and Hall study that was not included in model training (**Fig. 4**) (21). The predictions showed sufficient agreement with the data (39.6 < 265 = ℱ^cdf−inv^(0.95,229)). Both predictions and experimental data showed rapid increases in total body mass (**Fig. 4A**) and fat mass (**Fig. 4B**) following *ad libitum* access to the high-fat diet (**Fig. 4**, black bar), with the opposite pattern observed after switching back to chow. The model also predicted energy intake with sufficient agreement throughout the diet-switching intervention (**Fig. 4C**).

**Figure 4.**
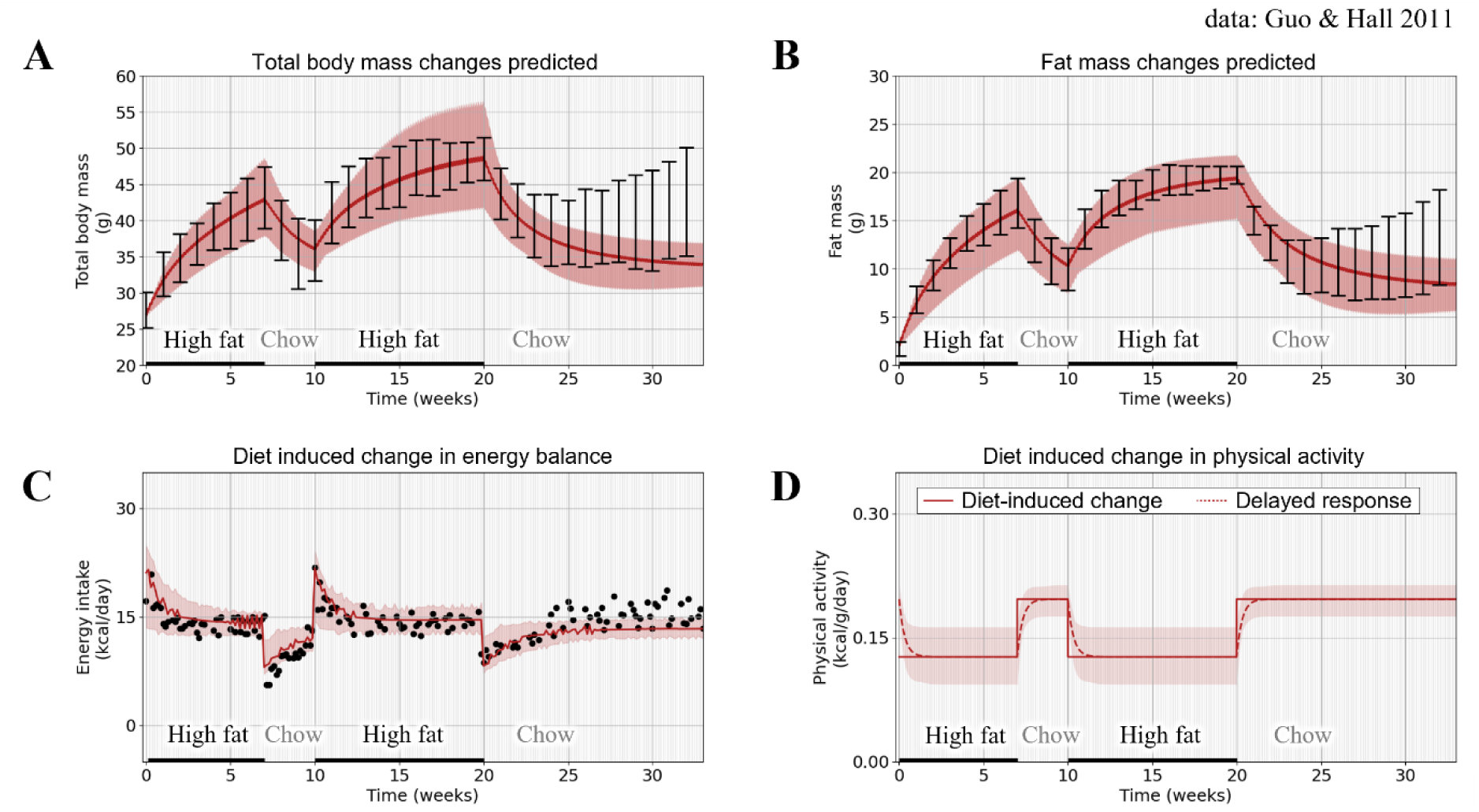
Model predictions of a mouse study (21) switching between a high-fat diet and a chow diet. Lines represent the prediction obtained using the best fit to all estimation data jointly (Figs. 2–3 and **Figs. S1–S2**), coloured areas indicate model uncertainty, grey areas indicate night-time, and error bars show population data (mean ± SEM). **(A)** Total body mass data and model prediction. **(B)** Fat mass data and model prediction. **(C)** Energy intake data and model prediction. **(D)** Predicted diet-induced changes in energy expenditure from physical activity. Energy expenditure from physical activity is described as a delayed response (dotted line and shaded area) to the diet-induced change (solid line).

The physical activity parameter was assumed to equal the value estimated for mice consuming the same diet in the estimation dataset (**Fig. 4D**). The best agreement with the data was obtained when physical activity was higher during chow feeding than during high-fat feeding (**Fig. 4D**, line), consistent with the original study (21). Acceptable agreement was also achieved with similar activity levels across diets (**Fig. 4D**, shaded area), suggesting that differences between diets could partly be explained by differences in energy intake (**Fig. 4C**).

### Model validation: Predicting the effect of the GLP-1RA exenatide on eating behaviour

Model validation was continued by evaluating the model’s ability to describe and predict the effect of the GLP-1RA exenatide on appetite using two pre-existing studies (**Fig. 5**).

**Figure 5.**
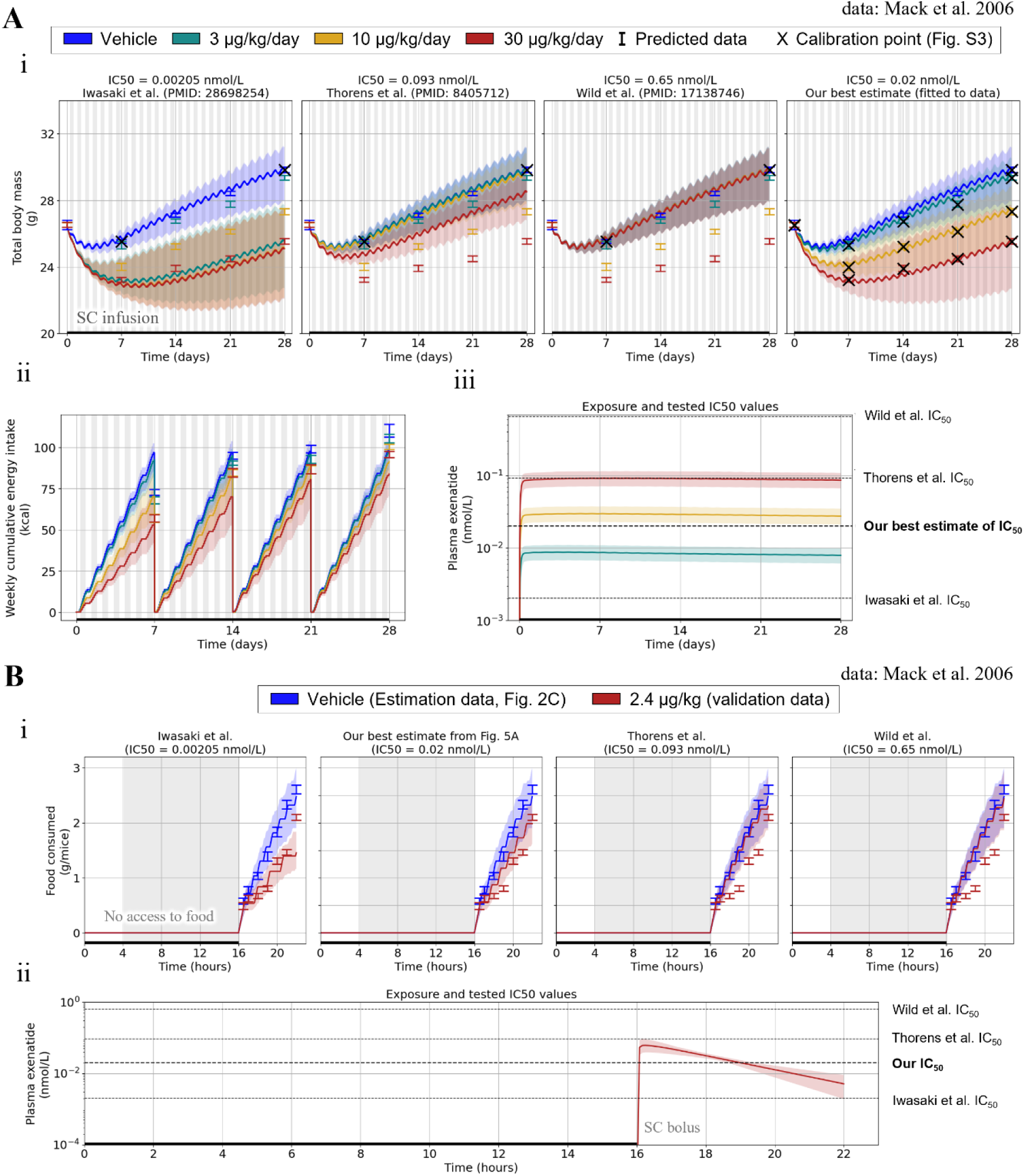
Model predictions of exenatide effects on body mass and energy intake. Lines represent the prediction obtained using the best fit to all estimation data jointly (Figs. 2–3 and **Figs. S1–S2**), grey areas indicate night-time, coloured areas indicate model uncertainty, and error bars show population data (mean ± SEM). **(A)** Model predictions of an exenatide mouse study (28) consuming a high-fat diet (D12331), separated into four groups: vehicle (blue), 3 µg/kg/day subcutaneous (SC) infusion (turquoise), 10 µg/kg/day SC infusion (yellow), and 30 µg/kg/day SC infusion (red). **(i)** Four model predictions of total body mass changes using three reported exenatide IC_50_ values (34–36) and an IC_50_ value estimated from the data using the model. Data marked with an “x” were used for calibration: the vehicle group was used to estimate energy expenditure from physical activity (**Fig. S3**), while the treated groups were used to estimate the IC_50_ used in the fourth prediction. **(ii)** Predicted energy intake using the calibrated IC_50_ value. **(iii)** Predicted exenatide exposure, with the tested IC_50_ values represented by horizontal lines. **(B)** Model simulations of a mouse study (28) in which the vehicle group (blue; estimation data) was used as estimation data (Fig. 2C) and the exenatide-treated group was predicted (red; validation data). Mice from both groups were refed chow diet (Teklad LM-485) after 16 h of fasting (black line on the x-axis). The study was predicted using the same pharmacodynamic parameters as in panel A. **(i)** Food consumption and model simulations. **(ii)** Predicted exenatide exposure, with the tested IC_50_ values represented by horizontal lines.

The first study reported total body mass (**Fig. 5A i**) and weekly cumulative energy intake (**Fig. 5A ii**) in mice with *ad libitum* access to high-fat diet (D12331) (28). All mice underwent subcutaneous implantation of an osmotic pump. Exenatide exposure was predicted using the GLP-1RA module (**Fig. S1**), with the highest dose reaching a steady-state plasma concentration of approximately 0.1 nmol/L (**Fig. 5A iii**). Naive (**Fig. S3 i**) and calibrated predictions (**Fig. S3 ii**) were generated by calibrating two free parameters to vehicle data (**Fig. S3 iii**). Physical activity in all treated groups was assumed to match the vehicle group (**Fig. 5A**, blue, marked with ‘x’). Using this vehicle-based calibration, the effect of exenatide was predicted (**Fig. 5A i**) using an assumed maximum inhibition of 75% (28), an assumed Hill coefficient fixed at 2 to represent a sigmoidal exposure–response relationship while reducing the number of pharmacodynamic parameters requiring estimation (32,33), and three different exenatide IC_50_ values reported in the literature: 0.00205 nmol/L (34), 0.093 nmol/L (35), and 0.75 nmol/L (36). Estimating the IC_50_ from the data yielded a best fit of 0.02 nmol/L (**Fig. 5A i**, right), which was then used to predict weekly cumulative energy intake (**Fig. 5A ii**). Three key observations were made. First, an initial stress response was required to describe the data, potentially reflecting surgery or adaptation to handling (**Fig. 5A i**, initial drop in mass; **Fig. S3A iii**, initial stress). This response was implemented as an effect on energy expenditure, although an effect on energy intake could not be excluded, as vehicle energy intake was overpredicted at the first measurement (**Fig. 5A ii**, 7 days, blue). Second, the model predicts a diminishing exenatide efficacy over time (**Fig. 5A ii**), which is explained in the model as mice gradually regulating their food intake based on energy intake. A diminishing effect may also be explained by changes in exposure from the osmotic pump (37,38). Third, the model showed sufficient agreement with the data 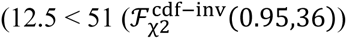.

In the second study, outcomes for an exenatide-treated group from the short-term Mack et al. study were predicted (28) (**Fig. 5B i**), with only the corresponding vehicle data used for model training (**Fig. 2C**). Exenatide exposure was predicted from the study-reported 2.4 µg/kg subcutaneous bolus (**Fig. 5B ii**). Of the four previously tested potency values, the IC_50_ value estimated from the month-long Mack et al. study showed the best agreement with data in Figure 5A 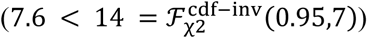. These results support cross-study validation across mouse strains (C57BL/6J to NIH Swiss), diets (high-fat to chow), and study durations (one month to 6 h of refeeding). The model also qualitatively agreed with literature reports that GLP-1RA treatment reduces meal frequency (39) and meal size (40), as reflected in the predicted two fewer meals and smaller first meal during refeeding (**Fig. 5B i**, 16 h).

### Model validation: Predicting the effect of the GLP-1RA semaglutide on eating behaviour

Model validation was continued by predicting the effect of the GLP-1RA semaglutide using one new and one pre-existing study (**Fig. 6**).

**Figure 6.**
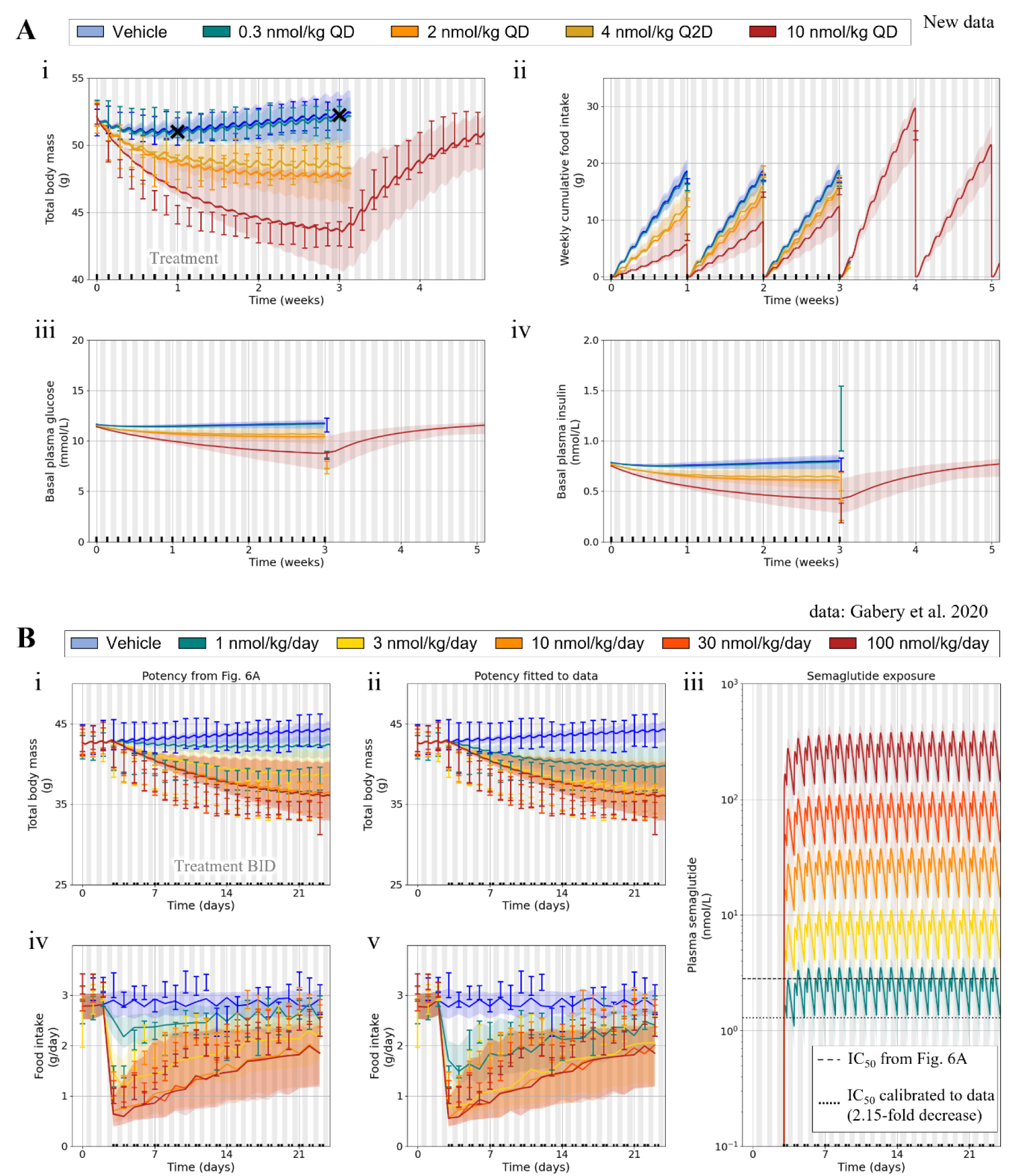
Model predictions of the effects of semaglutide on total body mass, energy intake, and disease prevention. Lines represent the prediction with the best fit to all estimation data jointly (Figs. 2–3 and **Figs. S1–S2**), coloured areas indicate model uncertainty, grey areas indicate night-time, and error bars show population data (mean ± SEM). **(A)** Model predictions of a new semaglutide study in diet-induced obesity (DIO) mice with ad libitum access to a high-fat diet (D12492). The mice were assigned to five groups: vehicle (blue), 0.3 nmol/kg/day QD subcutaneous (SC) injections (turquoise), 2 nmol/kg/day QD SC injections (orange), 4 nmol/kg/day Q2D SC injections (yellow), and 10 nmol/kg/day QD SC injections (red). Data marked with an “x” were used to estimate energy expenditure from physical activity (**Fig. S3**), while the treated groups were used to estimate the semaglutide pharmacodynamic parameters (IC_50_ and Imax). **(i)** Total body mass changes. **(ii)** Weekly cumulative food intake. **(iii)** Basal plasma glucose concentrations. **(iv)** Basal plasma insulin concentrations. **(B)** Model predictions of a semaglutide study in mice (3) with ad libitum access to a high-fat diet (D12451). The mice were assigned to six groups: vehicle (blue), 1 nmol/kg/day BID SC injections (turquoise), 3 nmol/kg/day BID SC injections (yellow), 10 nmol/kg/day BID SC injections (light orange), 30 nmol/kg/day BID SC injections (dark orange), and 100 nmol/kg/day BID SC injections (red). Semaglutide treatment started on day 3 (black line on the x-axis). **(i)** Total body mass predictions using the same parameters estimated from the study shown in Figure 6A. **(ii)** Total body mass predictions using a semaglutide IC_50_ value calibrated to the data. **(iii)** Predicted semaglutide plasma concentrations, with the tested IC_50_ values indicated as horizontal lines. **(iv)** Food intake predictions using the same parameters estimated from the study shown in Figure 6A. **(v)** Food intake predictions using a semaglutide IC_50_ value calibrated to the data.

In the new study, mice had *ad libitum* access to a high-fat diet (D12492) and were assigned to vehicle (**Fig. 6A**, blue) and four semaglutide-treated groups (**Fig. 6A**, non-blue). The model was fitted to endpoint exposure data and a short-term pharmacokinetic study simultaneously to predict the semaglutide exposure throughout the study (**Fig. S1C**). Physical activity was predicted and estimated as in the Mack et al. study (**Fig. 5A**), with a naive prediction (**Fig. S3B i**) followed by calibration to vehicle data (**Figs. S3B ii–iii**). The semaglutide pharmacodynamic parameters were estimated from the treated groups (Imax = 0.9; IC_50_ = 2.8 nmol/L), assuming a Hill coefficient of 2, as for exenatide. Model predictions of total body mass (**Fig. 6A i**), weekly cumulative food intake (**Fig. 6A ii**), basal plasma glucose (**Fig. 6A iii**), and basal plasma insulin (**Fig. 6A iv**) were compared with the data, and five key observations were made. First, the model captured day-to-day body mass fluctuations in the Q2D 4 nmol/kg and QD 2 nmol/kg groups, including increases on non-dosing days (**Fig. 6A i**, yellow and orange). Second, the model adequately predicts all groups (81 < 180.7 = ℱ^cdf−inv^(0.95,151)).

Third, body mass loss was underpredicted in the highest-dose group despite adequate weekly food intake predictions (**Fig. 6A**, red, week 1), potentially reflecting an initial stress response or time-dependent changes in energy expenditure not captured by the minimal calibration. Fourth, the model predicted an initial overshoot in food intake relative to baseline post semaglutide treatment (**Fig. 6A ii**, red, week 4), followed by a subsequent return towards baseline (**Fig. 6A i**, red). Finally, predicted changes in basal glucose and insulin were physiologically plausible (**Figs. 6A iii–iv**), with baseline values assumed to match the vehicle group. The glucose-insulin dynamics were predicted using the glucose-insulin module (**Fig. S2**).

The same pharmacodynamic parameters were used to predict an independent, pre-existing semaglutide study (3) with regard to total body mass (**Figs. 6B i–ii**) and food intake (**Figs. 6B iv–v**). Semaglutide was administered as a subcutaneous bolus twice daily and its exposure was predicted using the GLP-1RA module (**Fig. 6B iii**). The same approach to predicting and estimating physical activity was used in this study as described previously, with an initial naive prediction (**Fig. S3C i**), followed by calibration to vehicle data (**Figs. S3C ii–iii**). Body mass was well predicted except at the lowest dose, where the semaglutide effect was underpredicted (**Fig. 6B**, turquoise). A 2.15-fold reduction in IC_50_ improved agreement (**Fig. 6B v**). Because exposure was predicted rather than measured, this discrepancy may reflect exposure uncertainty rather than a true potency difference, as exposure differences have been observed in diseased mouse models (4,41). Nevertheless, both the uncalibrated (χ² = 205 < 328) and calibrated (χ² = 169 < 328) predictions agreed with the data, capturing body mass loss towards 36 g (**Figs. 6B i–ii**) and diminishing effects on food intake over time (**Figs. 6B iv–v**).

### Model utility: Connecting appetite-driven food intake to metabolic changes across diet and GLP-1RA treatments in mice

As a final model evaluation, the model was used to predict dietary and GLP-1RA treatment interventions in healthy and obese mice (**Fig. 7**). Five key model components provide mechanistic context for these predictions (**Figs. 7A**). First, physical activity-related energy expenditure decreases with high-fat feeding and higher body mass (**Fig. 7A i**), broadly consistent with literature observations (42–44). Second, appetite increases over the duration of studies (**Fig. 7A ii**). Third, appetite follows a sinusoidal circadian pattern with the lowest value at 2 pm and the highest at 2 am (**Fig. 7A iii**), resulting in more meals being consumed at night. Fourth, mice strongly self-regulate appetite according to energy intake (**Fig. 7A iv**). Finally, non-lipids contribute approximately 21-fold more to direct satiety per calorie than lipids (**Fig. 7A v**; uncertainty range: 5.5–57.9).

**Figure 7.**
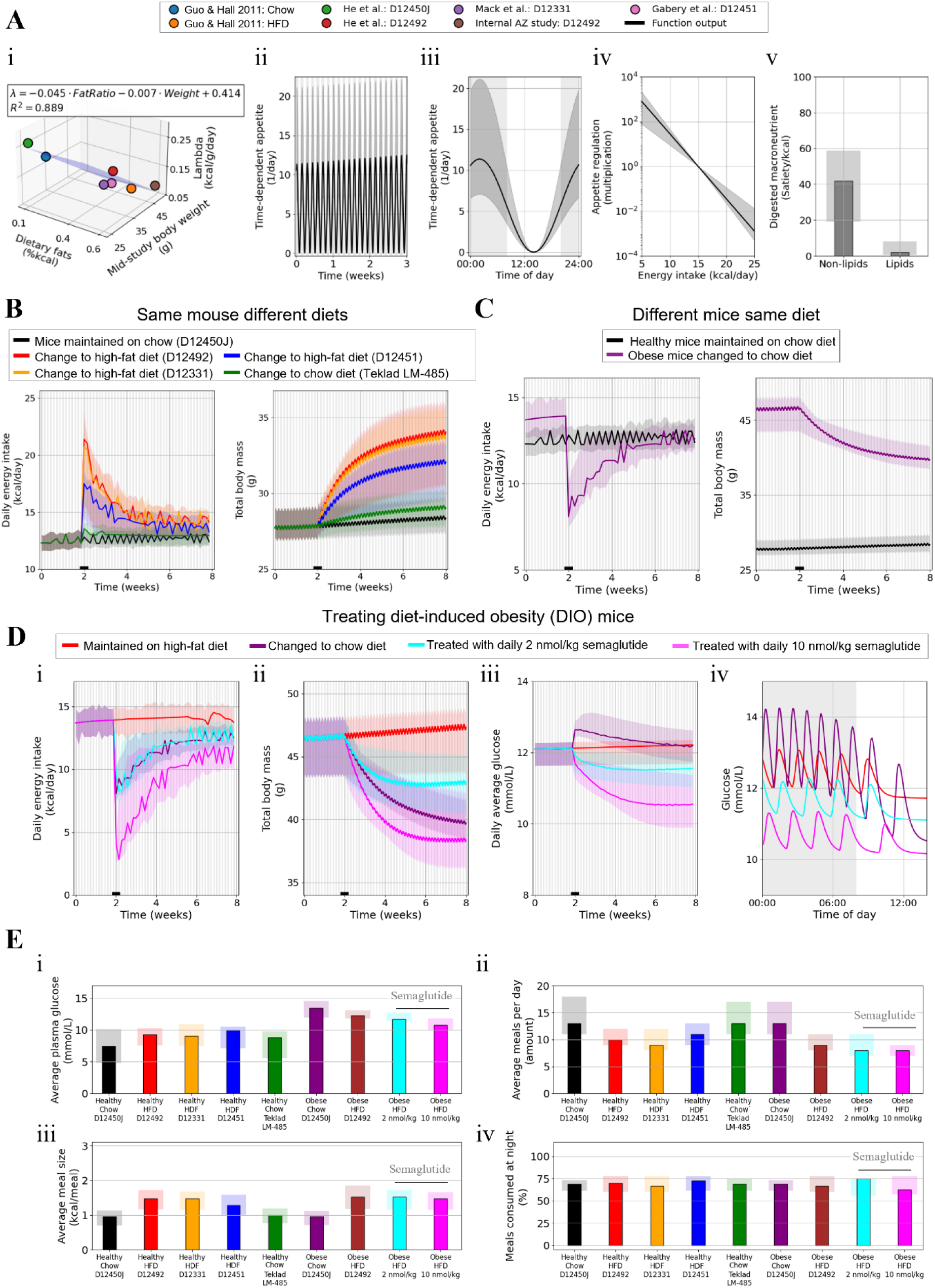
Model predictions of dietary and GLP-1RA effects on mouse appetite, body mass, and disease prevention. Lines and bars represent predictions with the best fit to all estimation data jointly (Figs. 2–3 and **Figs. S1–S2**) and the new semaglutide study (Fig. 6A). Coloured areas indicate model uncertainty, vertical grey areas indicate night-time, and black lines on the x-axis indicate simulation changes. **(A)** Predicted variables describing the model components regulating eating behaviour. **(i)** Predicted study-specific energy expenditure from physical activity relative to study-specific covariates. **(ii)** Increasing appetite development over time. **(iii)** Appetite development from circadian rhythm. **(iv)** Self-regulation of food intake according to energy intake. **(v)** Macronutrient-specific direct effect on satiety. **(B)** Simulated healthy mice switched to different diets. **(C)** Healthy and obese mice switched to the same diet. **(D)** Obese mice on a high-fat diet switched to a chow diet or treated with semaglutide. **(i)** Daily energy intake. **(ii)** Total body mass. **(iii)** Daily average plasma glucose concentrations. **(iv)** Plasma glucose concentrations on the final day of the simulation (day 56). **(E)** Comparison of the final day of all simulated interventions. **(i)** Average plasma glucose. **(ii)** Average number of meals. **(iii)** Average energy per meal. **(iv)** Proportion of total meals consumed at night.

The effects of five diets on energy intake and total body mass were compared (**Fig. 7B**). Mice were simulated on chow (D12450J) for 6 weeks before switching to alternative diets. Three key observations emerged. First, switching to high-fat diets caused an initial energy intake spike followed by a decline towards 14 kcal/day, attributed to differences in macronutrient-dependent satiety (**Fig. 7A v**) and caloric self-regulation (**Fig. 7A iv**). Second, the best agreement with data was achieved with incomplete compensation for diet-specific caloric density differences (**Fig. 7B**, lines), although sufficient agreement was also obtained with full compensation (**Fig. 7B**, shaded areas). Both scenarios produced diet-induced obesity through diet-dependent differences in energy expenditure (**Fig. 7A i**). Finally, all alternative diets increased body mass faster than continued chow feeding, although body mass slowly increased over time even with chow (**Fig. 7B**, black).

Healthy mice maintained on chow (D12450J) were compared with obese mice switched from high-fat diet (D12492) to the same diet as the healthy mouse (**Fig. 7C**). The obese mice were predicted to initially lose body mass rapidly, followed by a slower decline. By week 8, both groups consumed similar daily energy intakes, while healthy mice slowly gained body mass and obese mice continued to lose it.

Dietary and semaglutide treatments of obese mice were compared in terms of daily energy intake (**Fig. 7D i**), total body mass (**Fig. 7D ii**), daily average glucose (**Fig. 7D iii**), and glucose meal responses (**Fig. 7D iv**). In these simulations, obese mice were first maintained on a high-fat diet (D12492) for 6 weeks, followed by either a switch to chow (**Fig. 7D**, magenta) or received daily subcutaneous semaglutide boluses of 2 or 10 nmol/kg. Five key observations emerged. First, switching to chow produced a similar energy intake but faster body mass loss than 2 nmol/kg semaglutide **(Fig. 7D i–ii)**, attributed to higher physical activity-related energy expenditure (**Fig. 7A i**). Second, daily average glucose initially increased after the switch to chow, reflecting its higher carbohydrate content, before gradually declining as metabolic regulation improved (**Fig. 7D iii**).

Third, despite faster body mass loss, chow-fed mice showed less stable glucose meal responses than mice treated with 2 nmol/kg semaglutide, failing to return to baseline glucose overnight because of metabolic dysregulation, nocturnal eating habits, and higher carbohydrate intake (**Fig. 7D iv**). Fourth, 10 nmol/kg semaglutide produced the greatest reductions in energy intake, body mass, average glucose, and glucose meal responses (**Fig. 7D**, pink). Finally, the body mass difference between chow-fed and high-dose semaglutide-treated mice diminished (**Fig. 7D ii**). This is because energy expenditure in mice on the chow diet remains consistently higher, while the difference in energy intake decreases over time as the mice regulate their food intake.

Outcomes on the final simulation day were compared across interventions (**Fig. 7E**). Obese mice switched to chow were predicted to have the highest average plasma glucose (**Fig. 7E i**), reflecting metabolic dysregulation combined with higher carbohydrate intake. Mice on the chow diet are predicted to eat 12–18 meals per day and consume more frequent meals with lower caloric content compared with mice on the high-fat diet (**Figs. 7E ii-iii**), all of which is in qualitative agreement with the literature (45–47). For the semaglutide treatments, the model predicts reductions in average plasma glucose levels (**Fig. 7E i**), total energy intake (**Figs. 7E ii-iii**), and eating frequency (**Fig. 7E ii**), which is in general agreement with the literature (39,40,45). The fraction of meals consumed at night is predicted to remain unchanged across all interventions, ranging from 60% to 80% (**Fig. 7E iv**), which is in qualitative agreement with the literature (48,49).

Taken together, the model training (**Figs. 2-3** and **Figs. S1-S2**), validation (**Figs. 4-6**), and utility (**Fig. 7**) demonstrate a model capable of connecting appetite-driven food intake to metabolic changes across diet and GLP-1RA treatments in mice.

## DISCUSSION

In this study, we present and evaluate a mathematical model of mouse eating behaviour and the cumulative metabolic trajectory towards or away from obesity-associated diseases such as diabetes (**Fig. 1**). The strength of the model lies in its ability to predict appetite-driven meal patterns in response to experimental conditions, such as diet and GLP-1RA treatment, and to link these short-term eating behaviours to cumulative long-term changes in body mass and metabolic regulation. Using a minimal set of mechanisms, the model describes food intake across different dietary conditions (**Fig. 2**), changes in body mass and metabolic regulation (**Fig. 3**), and the effects of exenatide (**Fig. 5**) and semaglutide (**Fig. 6**) on appetite and downstream physiology. Model agreement with the data was evaluated using a data-driven hypothesis-testing approach in which all estimation data were considered jointly (**Figs. 2–3** and **Figs. S1–S2**), allowing the same model and parameter vectors to be evaluated across different mice, diets, treatment regimens, and experimental timescales (**Figs. 2-6**). The resulting model was then used to investigate dietary and GLP-1RA interventions, with predictions showing qualitative agreement with the literature (**Fig. 7**). Collectively, these findings demonstrate, to the best of our knowledge, a first mathematical model of mouse appetite that connects individual meal generation to the cumulative effects of diet and GLP-1RA treatment on eating behaviour, body mass, metabolic dysregulation, and glucose-insulin regulation.

Describing changes in mouse eating behaviour is critical when evaluating compounds that affect appetite, such as GLP-1RAs. Eating behaviour has often been summarised using daily or weekly energy intake (10,11), which presents two key shortcomings: it may obscure how a compound changes eating patterns, and it leads to a temporal conflict where energy intake averages are insufficient to capture the rapid effects of GLP-1RAs on individual meal responses. By explicitly modelling meal generation, we address both these shortcomings and reveal features of GLP-1RA treatment that can only be captured using a model describing both appetite-driven body mass changes and meal responses. For example, although diet-induced obese mice switching to a healthy chow diet were predicted to achieve a similar reduction in body mass after six weeks as mice treated with semaglutide, the predicted glucose dynamics differed substantially. Mice switched to chow initially showed an increase in daily average glucose, large glucose excursions following meals, and incomplete recovery towards baseline during the night after six weeks of treatment. In contrast, the semaglutide treatment rapidly reduced daily average glucose and resulted in improved glucose regulation and meal responses after the same treatment period. Importantly, the model goes beyond describing the rapid and slower effects of GLP-1RAs by also incorporating dietary effects, allowing both drug- and diet-related effects on eating habits to be distinguished (**Fig. 7**). This distinction provides a potential application in preclinical drug discovery, where understanding whether changes in energy intake arise from compound effects or dietary conditions can be important for interpreting experimental results.

Several assumptions underlying the modelling analysis should be considered. The appetite module is a semi-mechanistic and simplified representation of the physiological processes regulating eating behaviour and it is not a complete or definitive description. Alternative mechanisms may also explain the observed data, and the validity of the current model has only been tested within the interventions evaluated herein. Additional hypotheses were tested but not included in the model as they were not identifiable from the available data. These include lean- and fat-mass-dependent appetite development, carbohydrate- and protein-specific effects on satiety, between-mouse variability in circadian rhythm, expansion of the body mass module to describe faster-changing components such as water weight, age effects on appetite and energy expenditure, and a more complex GLP-1RA effect on satiety. We encourage future work to explore these hypotheses.

Beyond these structural assumptions, several practical limitations of the modelling framework could be addressed in future refinements. First, most of the data used to evaluate the model were available on a day-to-day basis, whereas minute-to-minute measurements would provide a stronger test of the mechanisms governing meal generation. Second, the digestive system was intentionally kept simplified and could be replaced by existing models with more detailed descriptions of gastric emptying and digestion (50–55). We chose not to incorporate these models to balance model complexity with the temporal resolution of the available data. Third, the GLP-1RA pharmacodynamic model does not describe receptor binding, receptor occupancy, or receptor turnover. Instead, a direct-response model using Imax, Hill coefficient, and IC_50_ values was used to describe the effects from GLP-1RA plasma exposure. Three previously reported exenatide IC_50_ values were evaluated, differing in their experimental origin and relevance to predicting *in vivo* effects in mice (**Fig. 5**): one value was estimated from food intake and body mass changes in C57BL/6J mice (34); another was estimated from *in vitro* receptor-transfected fibroblasts and GLP-1-induced cAMP production (35); and the third was obtained from an *in vitro* competitive receptor-binding study using Rip1Tag2 mouse insulinoma tissue (36). These values were tested to evaluate model predictions using minimal calibration data, and we do not claim that *in vitro* measurements can be directly used to predict *in vivo* studies without careful knowledge-driven calibrations (56,57). A fourth limitation is that GLP-1RA exposures had to be predicted for several studies because measured exposure was not reported, which may introduce discrepancies between predicted and true exposure. Within a given study, such discrepancies would affect the estimated IC_50_ consistently, but they may limit comparisons across studies. This was not apparent between the two exenatide studies (**Fig. 5**) but could contribute to the differences observed between the two semaglutide DIO studies (**Fig. 6**). Additional missing information also limited more detailed evaluation of the model. For example, the change in food intake observed in the Gabery et al. study (**Fig. 6B iv–v**, 10–13 h) may reflect events not documented in the available study reports, such as cage cleaning or mouse handling. Finally, estimation of energy expenditure from physical activity remains a key limitation. Physical activity is difficult to predict without explicit information about the study environment, and future work could investigate this using study-specific covariates (**Fig. S4**) or hierarchical population modelling.

Mathematical modelling has been used for many years to describe meal responses (58–60) and changes in body mass (21,22,24). We continue this tradition and extend its applications. A related mathematical model presented in 2021 generates meals based on a variable termed *physiological state* (61). This model describes human eating behaviour, including meal timing and size, using stochastic processes constrained by predefined habitual time points. It was subsequently extended to describe minute-to-minute food intake and long-term changes in body weight, plasma glucose, insulin sensitivity, energy expenditure, and glycaemic variability (62). This model provides an important basis for describing human eating behaviour and may support future translation of our proposed model to humans. Nonetheless, there are two key differences between our approaches. First, our approach uses a semi-mechanistic ODE-based model to predict eating behaviour from circadian rhythm, dietary composition, and GLP-1RA exposure rather than stochastic processes. Second, the scope and intended application differ. The model presented herein was specifically developed to describe mice with *ad libitum* access to one food option at a time, while the previous model describes humans in a different experimental setting. Our model also describes the effect of GLP-1RAs, while the human model does not. In this way, our model extends appetite modelling into a preclinical drug discovery setting and provides a platform for supporting mouse study design.

In conclusion, we have demonstrated a model capable of connecting food intake to metabolic changes across diet and GLP-1RA treatments in mice. We found that, to sufficiently describe the data, mice gradually adjusted their food intake substantially according to its energy content, and that non-lipids provide greater satiety per calorie consumed than lipids. The model can support mouse study design by predicting feeding behaviour under different experimental conditions and support preclinical drug discovery by separating dietary and GLP-1RA effects on appetite. More broadly, the model connects short-term changes in eating behaviour to the trajectory of metabolic dysregulation, supporting research into diet-induced metabolic diseases.

## METHODS

The mathematical modelling followed a data-driven hypothesis-testing approach described previously (63,64). The general methodology is briefly described below.

The mathematical model was constructed using ordinary differential equations (ODEs) and follows the same general formulation (**Eqs. 1A-C**).

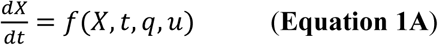

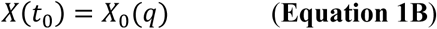

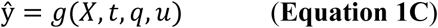

where *f* and *g* are smooth non-linear functions dependent on: *X* which is a vector of state variables, for example, lipids in the digestive system; *q* which is a vector of model parameters representing, for example, metabolic rates and scaling constants; *u* which is a vector of input signals, for example, drug administration or macronutrient profile of a diet; *X*(*t*_0_) which is a vector of initial conditions *X*_0_(*q*) that are defined as functions of the model parameters *q*; ŷ which is a vector of model outputs, for example, energy intake or body mass.

### Parameter estimation

The agreement between the model simulation and experimental data was evaluated using a weighted sum of squared residuals with a minimum error floor (**Eq. 2**):

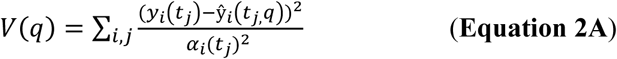

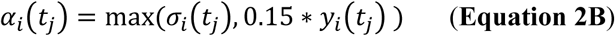

where *y_i_*(*t_j_*) is the measured mean of a group of data points; *j* is a specific data point within the experimental setup *i* at the time-point *t*; *σ_i_*(*t_j_*) is the standard error of the mean of the group; *α_i_* is the implemented minimum error floor; and ŷ*_i_*(*t_j_*,*q*) is the corresponding model-predicted value.

The weighted sum of squared residuals *V*(*q*) was minimised in an optimisation problem to find simulations with the best agreement with the data. For replicate-specific data points, such as the energy intake data used in Figure 2B i, the minimum error floor was increased from 15% to 35% to balance model fit between population- and individual-level data points.

Statistical *χ*^2^ tests were performed to evaluate model agreement with the data. The statistical test used the null hypothesis that the experimental data had been generated by the model, assuming that the experimental noise was additive, independent, and normally distributed (64). In practice, the agreement to data was rejected if the weighted sum of squared residuals *V*(*q*) (**Eq. 2A**) had a higher value than the *χ*^2^-threshold, given by the inverse *χ*^2^ cumulative density function (**Eq. 3**):

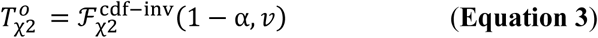

where 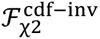 is the inverse cumulative distribution function; α is the significance level (α = 0.05 was used); and *v* is the degrees of freedom, which was equal to the number of data points in the dataset.

All mathematical modelling, simulations, plotting, and parameter estimation were performed in Python (Version 3.11) using the SUND Toolbox (65). The parameter estimation was executed in log-transformed parameter space using the particle swarm optimisation algorithm implemented in pyswarm. A swarm of 120 particles was initialised using a three-zone strategy where 30% of the particles were initialised from a local zone (±20%) from a manually determined initial guess, 35% from an intermediate zone (±200%), and 35% from a large zone (±600%). Optimisation was executed using eight parallel worker processes and continued for 200 iterations. Parameter uncertainty was assessed from all statistically non-rejected parameter vectors identified during the optimisation. Parameter vectors were accepted only if they were not rejected by the χ² test for any individual estimation study and satisfied the qualitative criteria. The qualitative criteria included: additional weighting of data points considered important for capturing qualitative behaviour, such as energy intake changes during dietary transitions; usage of condition-specific adjustments to minimise study-specific differences in energy expenditure from physical activity (66); and weights on model outputs to guide them towards physiologically sound magnitudes, such as a meal consumption rate and number of meals per day.

### Model calibrations of physical activity

To simulate body mass changes, energy expenditure from physical activity must be known or assumed. For studies included in the estimation data, physical activity was estimated from the data (**Figs. 2–3**).

For studies not included in the estimation data (**Figs. 5–6**), naive predictions were generated, assuming the same physical activity as in the estimation study with the most similar macronutrient profile to the predicted diet (**Fig. S3 A-C i**). These predictions were then complemented by estimating physical activity from vehicle data (**Figs. S3A–C ii**). This estimation included a study-start stress response (**Figs. S3A–C ii**, marked with ‘x’) and steady-state physical activity (**Figs. S3A–C ii**, marked with ‘+’). The resulting study-specific physical activity estimates are provided in the Supplementary Material (**Figs. S3A–C iii**) and compared with the physical activity assumed in the naive predictions (dotted black lines). The adaptation rate from the study-start stress response to steady-state physical activity was assumed to match the best fit to the estimation data (**Fig. S3A iii**, dotted blue line). In all studies, treated groups were assumed to have the same physical activity as the corresponding vehicle group (**Figs. 5–6**).

For simulations without data, vehicle-based calibration was not possible (**Fig. 7**). Instead, physical activity was predicted using information from all studies considered herein (**Fig. S4**). This was necessary to provide physiologically relevant predictions, as physical activity affects metabolic outcomes and may depend on mouse covariates, such as body mass, and dietary covariates, such as macronutrient profile. To investigate these relationships, study-specific covariates were compared with predicted study-specific physical activity using 16 regression models (**Fig. S4**). Of these models, a linear regression model with two predictors was selected for the simulations without calibration data, to balance low residuals (R^2^ = 0.889) with model simplicity (**Fig. 7A i**). This regression model predicts lower energy expenditure from physical activity in mice with higher body mass and those consuming a high-fat diet, broadly consistent with observations in the literature (42–44).

### Mathematical model structure

The mathematical model of appetite consists of eight optimised model parameters and six ODEs: one state describing appetite, one describing a delayed energy intake signal used for self-regulation of appetite in mice, and four describing the digestive system. The initial values for the states were simulated using a 28-day steady-state simulation starting at midnight.

The first ODE described the state variable *Appetite* (**Eq. 4**).

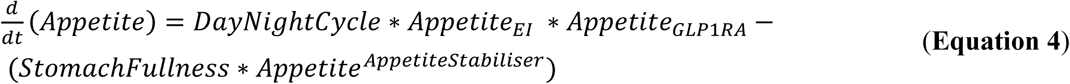

where *Appetite* is the model state describing the drive to eat in mice; *DayNightCycle* is the model function regulating appetite development according to the circadian rhythm (**Eq. 5**); *Appetite_EI_* is the model function regulating appetite development according to the energy intake (**Eq. 6**); *Appetite_GLP_*_1*RA*_ is the model function regulating appetite development due to GLP-1RA exposure (**Eq. 7**); *StomachFullness* is the model function regulating appetite development due to consumed macronutrients (**Eq. 8**); and *AppetiteStabiliser* is a model parameter stabilising appetite around the threshold generating meals (termed HungerThreshold in **Eq. 10**).

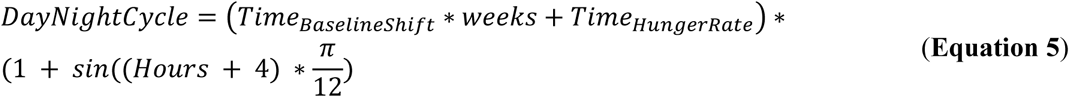

where *Time_BaselineS_*_ℎ*ift*_is a model parameter increasing the amplitude of *DayNightCycle* with study duration; *Time_HungerRate_* is a model parameter determining the amplitude of *DayNightCycle* at study start; *Hours* and *weeks* denote time elapsed since midnight on the first simulation day; and 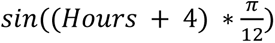 defines the sinusoidal function with a peak at 02:00.

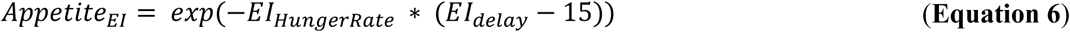

where *EI_HungerRate_* is a model parameter determining the magnitude of the fed state effect on appetite, 15 is a normalisation point of energy intake expressed as kcal/day, and *EI_delay_* is a delayed signal of energy intake expressed with an ODE (**Eq. 9**).

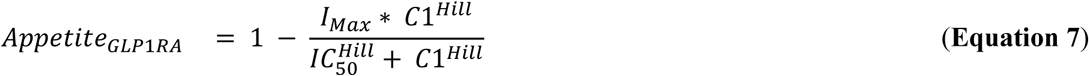

where *I_Max_* is the maximum GLP-1RA effect on appetite; *C*1 is GLP-1RA concentration in plasma; *Hill* is the Hill coefficient, which was assumed to be 2 for both semaglutide and exenatide; and *IC*_50_ is the plasma concentration yielding half of the maximum inhibition on appetite.

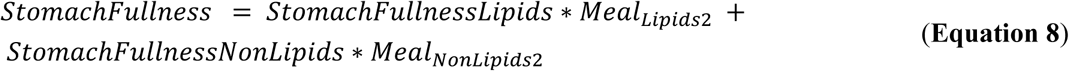

where *StomachFullnessLipids* is a model parameter determining the direct satiety effect per kilocalorie of lipids, while *StomachFullnessNonLipids* determines it for non-lipids; and *Meal_Lipids_*_2_ is the second digestive state for lipids (**Eq. 12**), while *Meal_NonLipids_*_2_ is the corresponding second digestive state for non-lipids (**Eq. 14**).

The previously described model function that regulates appetite based on the mice’s feeding history (**Eq. 6**) is dependent on a delayed state of energy intake (**Eq. 9**).

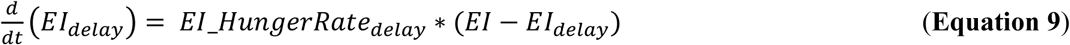

where *EI*_*HungerRate_delay_* is a model parameter describing the rate of the delay between *EI_delay_* and *EI*, which is the equation describing energy intake (**Eq. 10**).

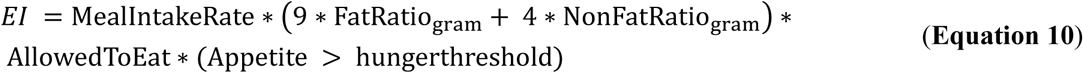

where 9 and 4 are the Atwater factor scaling of lipids and non-lipids from g to kcal (67); LipidRatio_gram_ is the percentage in mass of the diet consisting of lipids, and NonLipidRatio_gram_ is the same for non-lipids; MealIntakeRate is a model parameter describing meal consumption rate; AllowedToEat is a model input used to declare whether the mice have access to food (implemented as a one or zero); and (Appetite > hungerthreshold) is the Boolean condition that initiates and terminates meal consumption.

When a meal is consumed, the ingested food enters the model’s digestive system, which comprises four ODEs (**Eqs. 11-14**).

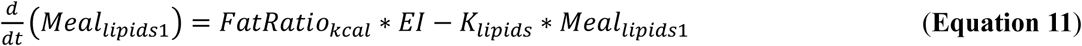

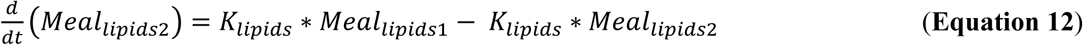

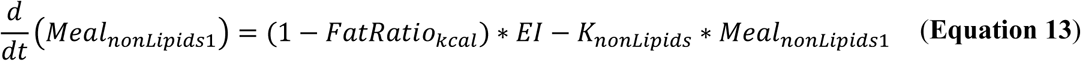

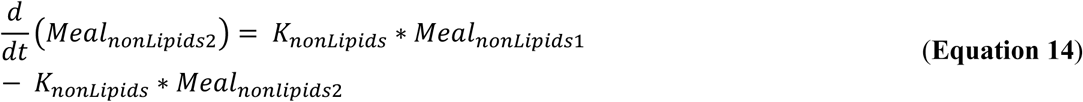

where *FatRatio_kcal_* is a model input representing fraction of dietary energy derived from lipids; *K_lipids_* is a lipid digestion rate fixed at 20 (1/day) based on gastric emptying rates in mice (68); and *K_nonLipids_* is a non-lipid digestion rate fixed at 60 (1/day) based on gastric emptying rates in mice (69,70) and consistent with a faster emptying rate than high-fat-containing meals in mice (68).

The final equation in the appetite module defines the carbohydrate input passed to the glucose–insulin module (**Eq. 15**).

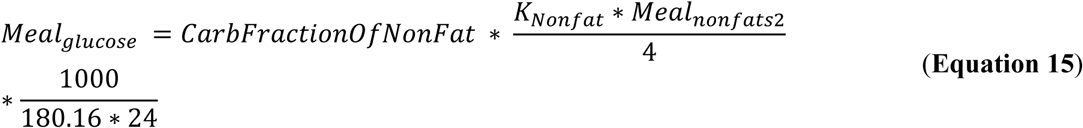

where *CarbFractionOfNonFat* is a model input representing the carbohydrate fraction from the non-lipid content in the diet; *K_Nonfat_* ∗ *Meal_nonfats_*_2_ is the emptying of non-lipids from the digestive system; 4 is the Atwater factor of carbohydrates, converting kcal to gram; and 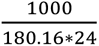 converts the unit from g/day to mmol/h.

Parameter values and uncertainty intervals are given in Table 1.

**Table 1.** Model parameters for the appetite module.

| **Name** | **Fitted value**<br>best (min-max) | **Unit** |
|---|---|---|
| *StomachFullnessNonLipids* | 41.93 (17.11- 91.10) | kcal<sup>-1</sup> |
| *StomachFullnessLipids* | 1.97 (0.70-11.04) | kcal<sup>-1</sup> |
| *Time*<sub>*HungerRate*</sub> | 5.67 (2.08-30.48) | day<sup>-1</sup> |
| *Time*<sub>*BaselineShift*</sub> | 0.19 (0.05 - 1.00) | 1/(day·week) |
| *EI*<sub>*HungerRate*</sub> | 0.67 (0.28-0.94) | day/kcal |
| *EI_HungerRate*<sub>*delay*</sub> | 0.02 (0.01-0.12) | day<sup>-1</sup> |
| *AppetiteStabiliser* | 1.79 (1.50-2.49) | dimensionless |
| MealIntakeRate | 16.87 (10.94 - 20.00) | g/day |

A detailed description of the three pre-existing models and how they were altered to support model-to-model crosstalk and their use in the new situations considered in this study is provided in the supplementary material (**Note S1**).

### Study design

The presented semaglutide study assessed the dose–response relationship of semaglutide on body weight and food intake, as well as post-withdrawal recovery in diet-induced obese mice.

All procedures were approved by the Gothenburg Ethics Review Committee on Animal Experiments (application 2002-2019). Male C57BL/6NTac diet-induced obese mice (Taconic, Denmark) were maintained under controlled conditions with *ad libitum* access to tap water and a high-fat diet (D12492; Research Diets, New Brunswick, NJ, USA). At 31 weeks of age, mice were randomised by body weight to vehicle (PBS, n = 10) or semaglutide at 0.3 or 2 nmol/kg once daily, 4 nmol/kg every other day, or 10 nmol/kg once daily (n = 8 per semaglutide group). Two 10 nmol/kg cohorts were included: one for terminal sampling at steady-state and one for post-treatment recovery. Treatments were administered subcutaneously at 10:00 for 22 days. Body weight was recorded daily and food intake weekly. After the final dose on day 21, all mice except those in the recovery cohort were fasted and euthanized under isoflurane for terminal blood collection. Insulin and glucose were measured by immunoassay (K152BZC-1; Meso Scale Discovery, Rockville, MD, USA) and enzymatic colorimetry on the ABX

Pentra 400 (Glucose HK CP, A11A01667, Horiba ABX, France), respectively. Semaglutide exposure was measured by LC-MS/MS. Exposure in the 0.3 nmol/kg once-daily group was below the lower limit of quantification (0.00101 µM) and was omitted from Figure S1C ii. The 10 nmol/kg once-daily recovery cohort was followed through day 33. On day 33, mice received a single 10 nmol/kg subcutaneous dose of semaglutide, and blood was sampled at 30 min and 1, 2, 4, 7, and 9 h post-dose for pharmacokinetic profiling (**Fig. S1C i**). Replicate- and population-level data are provided in the supplementary material (**Supplementary Table S1)**.

## DATA AVAILABILITY

All data used in this analysis are provided in the supplementary material and can also be downloaded from our GitHub repository (https://github.com/OscarSilfvergren/Connecting-Appetite-Driven-Food-Intake-to-Metabolic-Changes-in-Mice.git). Data collected from the literature were digitised using a publicly available software (71).

## CODE AVAILABILITY

All code used in the analysis is available from our GitHub repository (https://github.com/OscarSilfvergren/Connecting-Appetite-Driven-Food-Intake-to-Metabolic-Changes-in-Mice.git).

## ACKNOWLEDGEMENTS

We thank Henrik Podéus, Gustav Magnusson, Sasan Keshmiri, Kajsa Tunedal, Nicolas Sundqvist, William Lövfors and the other members of the team for developing and supporting the SUND Toolbox (65).

## FUNDING

Swedish Research Council 2023–03186 (GC)

Swedish Research Council 2023–05460 (GC)

X-HiDE, Knowledge Foundation 20200017 (GC)

CENIIT 15.09 (GC)

The Horizon Europe project STRATIF-AI 101080875 (GC)

The Swedish Fund for Research without Animal Experiments F2019-0010 (GC)

ELLIIT 2020-A12 (GC)

The County Council of Östergötland RÖ-1001928 (GC)

VINNOVA VisualSweden 2020-04711 (GC)

## AUTHOR INFORMATION

### Authors and Affiliations

**Linköping University, Linköping, Sweden**

Oscar Silfvergren, Peter Gennemark, and Gunnar Cedersund

**SUND sound medical decisions, Linköping, Sweden**

Oscar Silfvergren and Gunnar Cedersund

**AstraZeneca, Gothenburg, Sweden**

Christian Simonsson, Monika Sundqvist, Pär Nordell, and Peter Gennemark

**Örebro University, Örebro, Sweden**

Gunnar Cedersund

### Contributions

**Conceptualization:** OS, CS, PG, GC

**Methodology**: OS, CS, MS, PG, GC

**Investigation**: OS, CS, MS

**Visualization**: OS, CS, MS, PN, PG, GC

**Funding acquisition**: PN, GC

**Project administration**: MS, PN, GC

**Supervision**: PG, GC

**Writing – original draft**: OS

**Writing – review & editing**: OS, CS, MS, PN, PG, GC

### Corresponding author

Correspondence to Gunnar Cedersund.

## COMPETING INTERESTS

Christian Simonsson, Monika Sundqvist, Pär Nordell, and Peter Gennemark are employees of AstraZeneca and hold shares and/or stock options. Gunnar Cedersund is founder and CEO of SUND sound medical decisions, which commercialises digital twins and data-driven hypothesis testing. Oscar Silfvergren is an employee of SUND sound medical decisions.

## SUPPLEMENTARY MATERIALS

**Supplementary Table S1:** Data from the new semaglutide study

**Supplementary Table S2–S9**: All digitised data used in this study.

**Supplementary Note S1**: Detailed description of the body mass, glucose–insulin, and GLP-1RA modules

**Figure S1.**
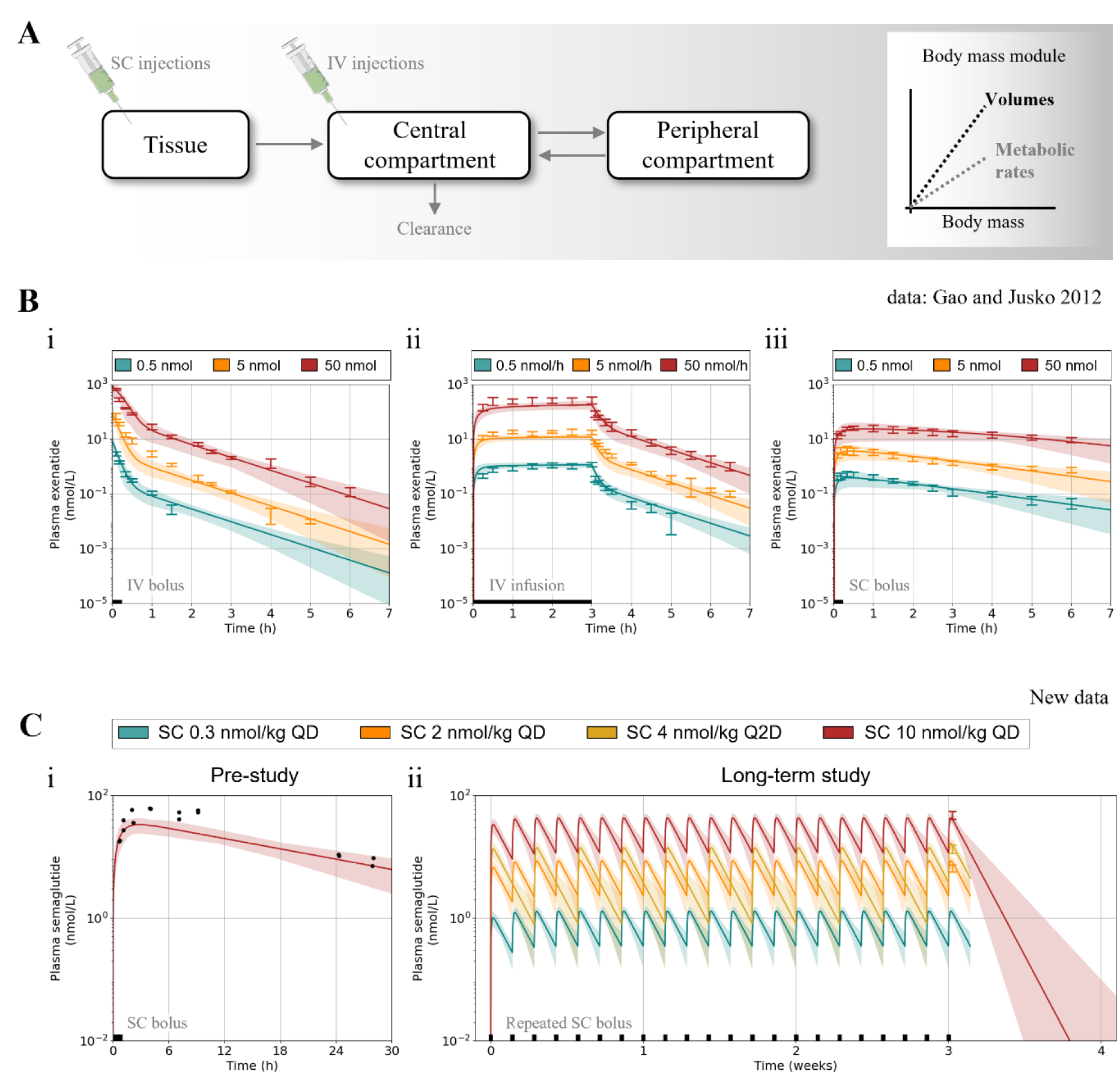
The GLP-1RA pharmacokinetic module used to describe exenatide and semaglutide exposures. Lines represent the best fit of the model to all estimation data jointly (**Figs. 2-3** and **Figs. S1-S2**), coloured areas indicate model uncertainty, and error bars show population data (mean ± SEM). **(A)** Illustration of the GLP-1RA module. **(B)** Model agreement with an exenatide rat study (72). The black line on the x-axis indicates the administration time. **(i)** Three intravenous (IV) bolus: 0.5 nmol (turquoise), 5 nmol (yellow), and 50 nmol (red). (**ii)** Three IV infusions: 0.5 nmol/h (turquoise), 5 nmol/h (yellow), and 50 nmol/h (red). **(iii)** Three subcutaneous (SC) injections: 0.5 nmol (turquoise), 5 nmol (yellow), and 50 nmol (red). **(C)** Model agreement with two new semaglutide studies in mice. The black line on the x-axis indicates the administration time. **(i)** Model agreement with data following a single 10 nmol/kg SC injection. **(ii)** Model agreement with data from four SC dosing regimens: 0.3 nmol/kg once daily (quaque die, QD) (blue), 2 nmol/kg QD (green), 4 nmol/kg every other day (quaque altera die, Q2D) (yellow), and 10 nmol/kg QD (red).

**Figure S2.**
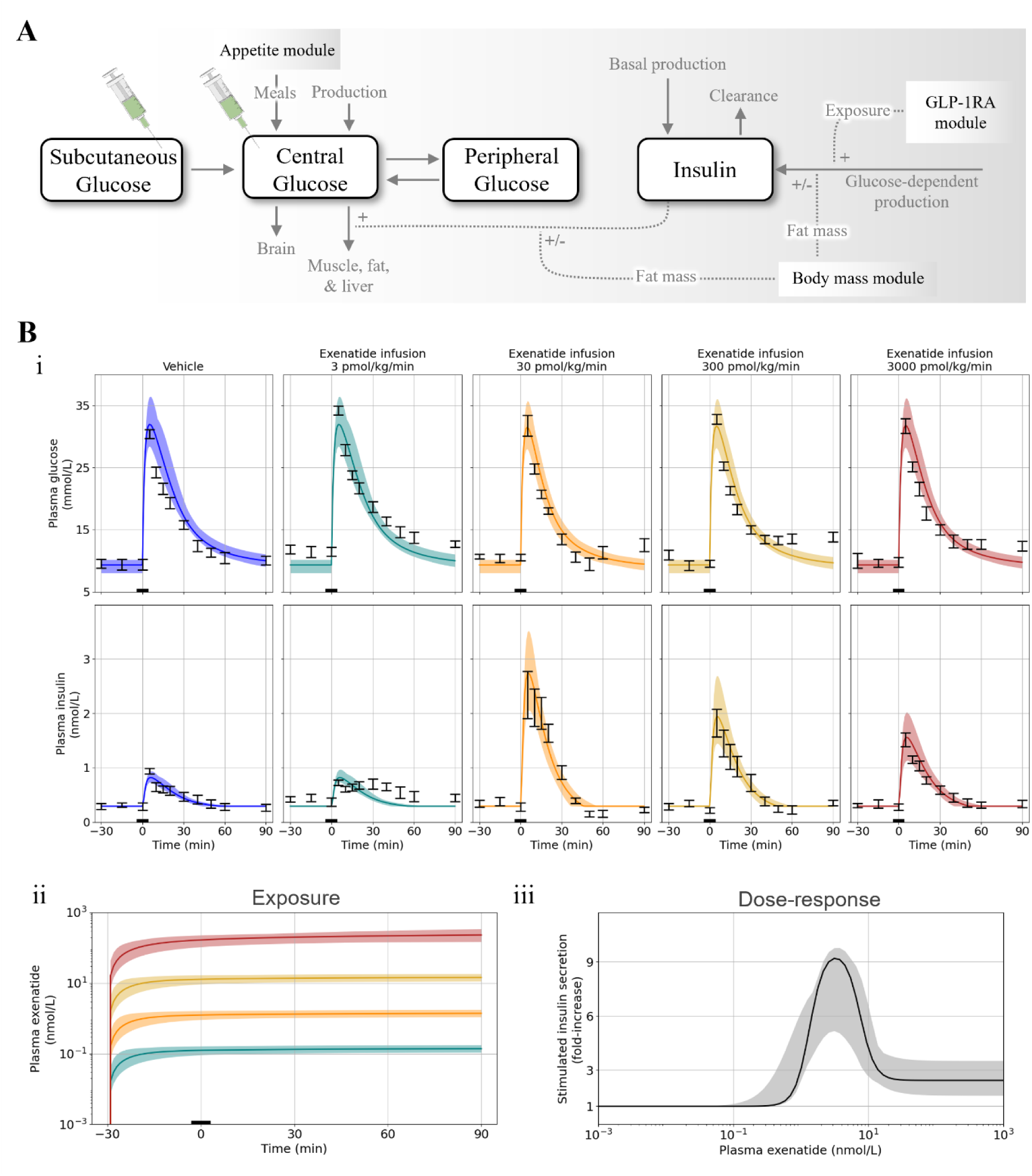
The glucose-insulin module and its fit to rat glucose injection data at various exenatide exposures. Lines represent the best fit of the model to all estimation data jointly (**Figs. 2-3** and **Figs. S1-S2**), coloured areas indicate model uncertainty, and error bars show population data (mean ± SEM). **(A)** Illustration of the glucose-insulin module and its crosstalk with the other three modules. **(B)** Model agreement with a rat study (72) comprising five groups: vehicle (blue), 3 pmol/kg/min intravenous (IV) infusion (turquoise), 30 pmol/kg/min IV infusion (yellow), 300 pmol/kg/min IV infusion (gold), and 3000 pmol/kg/min IV infusion (red). Black line on the x-axis indicates a glucose injection (5.7 mmol/kg) administered 30 min after the start of the exenatide IV infusions. **(i)** Data and model predictions of plasma glucose and plasma insulin. **(ii)** Predicted exenatide exposure using the GLP-1RA pharmacokinetic module (**Fig. S1**) and the exenatide exposure predicted to produce the maximum effect (EC_100_) on insulin secretion (dotted line). (**iii) The** predicted effect of exenatide on stimulated insulin secretion at various plasma exposures.

**Figure S3.**
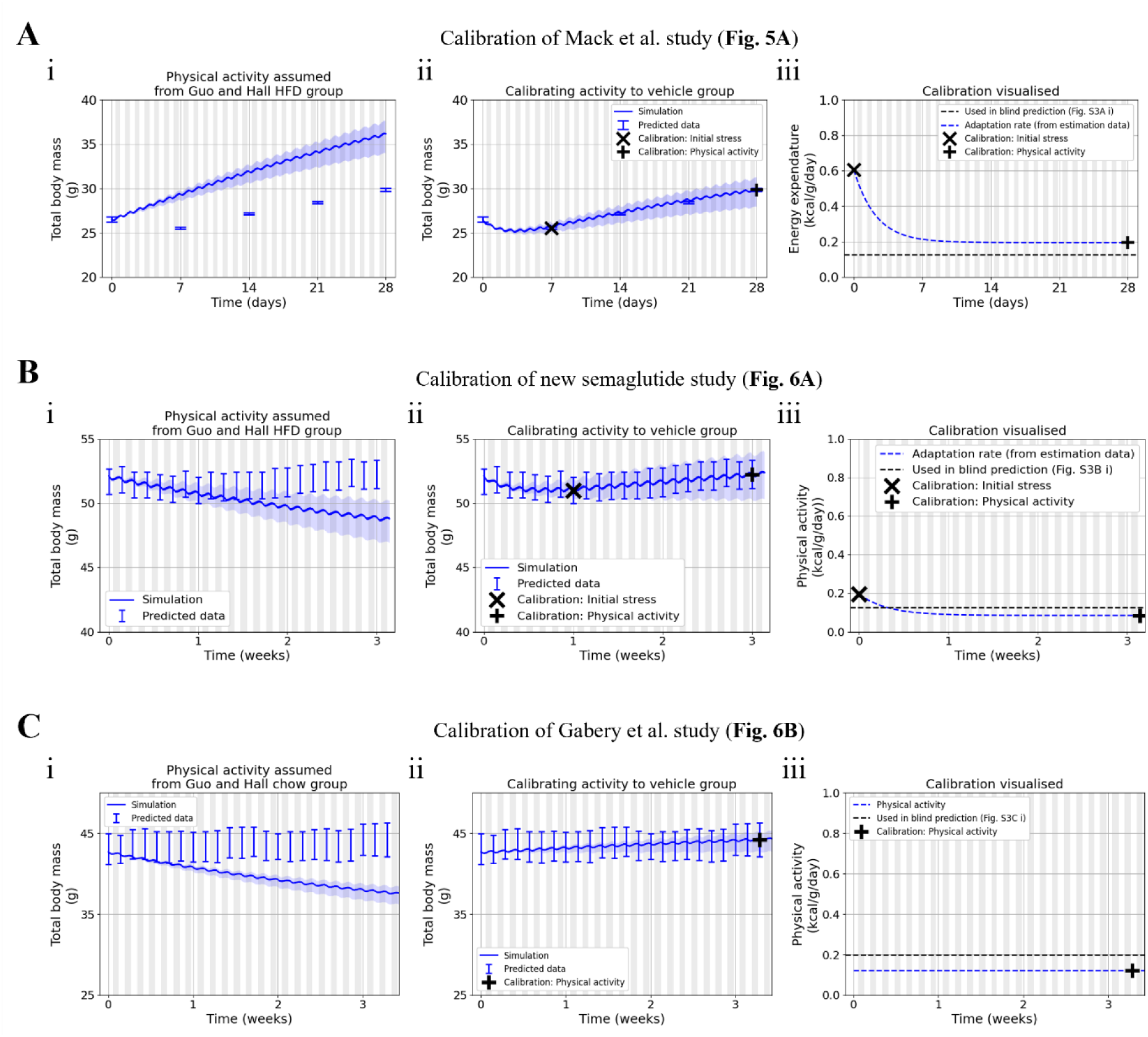
Calibrations used in the manuscript. Lines represent the prediction with the best fit to all estimation data jointly (**Figs. 2-3** and **Figs. S1-S2**), coloured areas indicate model uncertainty, and error bars show population data (mean ± SEM). **(A)** Calibration of physical activity to the Mack et al. study (28) (**Fig. 5A**). **(i)** Prediction using physical activity assumed from the high-fat-diet group in the Guo and Hall study (21) **(Fig. 3B). (ii)** Prediction using calibrated physical activity based on two points: an initial stress represented by an increased physical activity (data marked with “**x**”) and steady-state physical activity (data marked with “**+**”). (**iii)** Resulting calibrated physical-activity profile. **(B)** Calibration of physical activity to the new semaglutide study (**Fig. 6A**). **(i)** Prediction using physical activity assumed from the high-fat-diet group in the Guo and Hall study (21) **(Fig. 3B). (ii)** Prediction using calibrated physical activity based on two points: an initial stress represented by increased physical activity (data marked with “**x**”) and a steady state-physical activity (data marked with “**+**”). (**iii)** Resulting calibrated physical-activity profile. **(C)** Calibration of physical activity to the Gabery et al. study (3) (**Fig. 6B**). **(i)** Prediction using physical activity assumed from the chow-diet group in the Guo and Hall study (21) **(Fig. 3B). (ii)** Prediction using calibrated steady-state physical activity (data marked with “**x**”). (**iii)** Resulting calibrated physical-activity profile.

**Figure S4.**
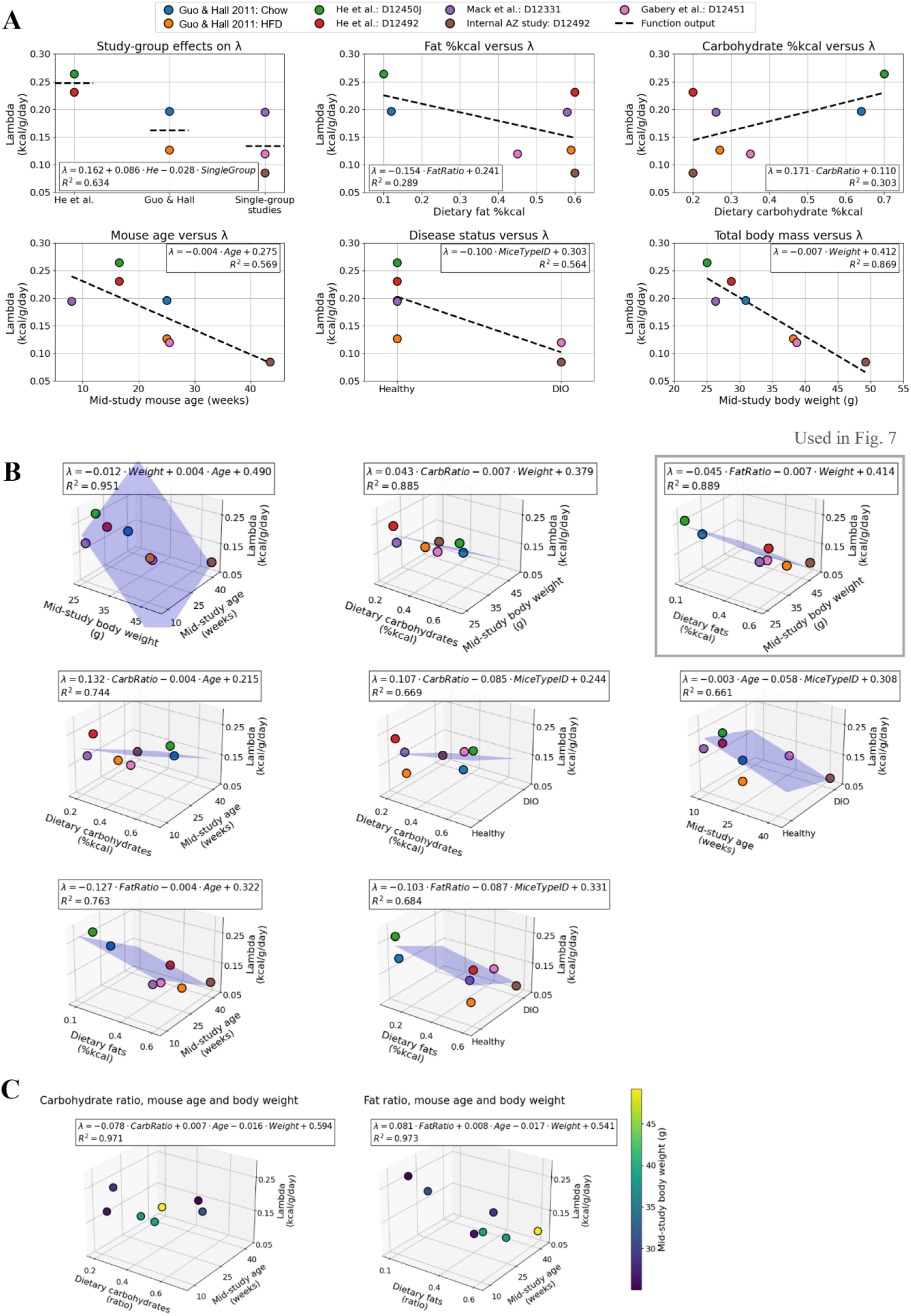
Connecting predicted study-specific physical activity with study-specific covariates. A total of seven different mouse groups from different studies were evaluated, for which data were available to estimate physical activity: two groups in the Guo and Hall study (21), one consuming chow (blue) and one consuming high-fat diet (orange) (the corresponding Research Diets product numbers not found in study report); two groups in the He et al. study (29), one consuming the chow diet (D12450J; green) and one consuming the high-fat diet (D12492; red); one group consuming the high-fat diet (D12331; magenta)in the Mack et al. study (28); one group consuming the high-fat diet (D12492; brown) in the new semaglutide study; and one group consuming the high-fat diet (D12451; pink) in the Gabery et al. study (3). The predicted physical activity of these seven mouse groups was compared with study-specific covariates. **(A)** Regression based on one predictor. **(B)** Regression based on two predictors. **(C)** Regression based on three predictors.

## Notes

https://github.com/OscarSilfvergren/Connecting-Appetite-Driven-Food-Intake-to-Metabolic-Changes-in-Mice.git

## References

1. Huang H, Zhao W, Qin N, Duan X. Recent Progress on Physiologically Based Pharmacokinetic (PBPK) Model: A Review Based on Bibliometrics. Toxics. 2024 Jun 14;12(6):433. doi:10.3390/toxics12060433 PubMed PMID: 38922113; PubMed Central PMCID: PMC11209072.

2. Loewa A, Feng JJ, Hedtrich S. Human disease models in drug development. Nat Rev Bioeng. 2023 Aug;1(8):545–59. doi:10.1038/s44222-023-00063-3

3. Gabery S, Salinas CG, Paulsen SJ, Ahnfelt-Rønne J, Alanentalo T, Baquero AF, et al. Semaglutide lowers body weight in rodents via distributed neural pathways. JCI Insight. 2020 Mar 26;5(6):e133429, 133429. doi:10.1172/jci.insight.133429 PubMed PMID: 32213703; PubMed Central PMCID: PMC7213778.

4. Bondarenko V, Løkke CR, Dobrowolski P, Mentzel CJ, Castro-Mejía JL, Hansen CHF, et al. Controlling the uncontrolled variation in the diet induced obese mouse by microbiomic characterization. Sci Rep. 2022 Aug 12;12:13767. doi:10.1038/s41598-022-17242-8 PubMed PMID: 35962158; PubMed Central PMCID: PMC9374709.

5. Barrett P, Mercer JG, Morgan PJ. Preclinical models for obesity research. Dis Model Mech. 2016 Nov 1;9(11):1245–55. doi:10.1242/dmm.026443 PubMed PMID: 27821603; PubMed Central PMCID: PMC5117231.

6. Young KG, McInnes EH, Massey RJ, Kahkoska AR, Pilla SJ, Raghavan S, et al. Treatment effect heterogeneity following type 2 diabetes treatment with GLP1-receptor agonists and SGLT2-inhibitors: a systematic review. Commun Med. 2023 Oct 5;3:131. doi:10.1038/s43856-023-00359-w PubMed PMID: 37794166; PubMed Central PMCID: PMC10551026.

7. Simonsen L, Lau J, Kruse T, Guo T, McGuire J, Jeppesen JF, et al. Preclinical evaluation of a protracted GLP-1/glucagon receptor co-agonist: Translational difficulties and pitfalls. PloS One. 2022;17(3):e0264974. doi:10.1371/journal.pone.0264974 PubMed PMID: 35245328; PubMed Central PMCID: PMC8896685.

8. Furió-Novejarque C, Díez JL, Bondia J. GLP-1 Receptor Agonists Models for Type 1 Diabetes: A Narrative Review. J Diabetes Sci Technol. 2025 Mar;19(2):332–9. doi:10.1177/19322968241285925 PubMed PMID: 39411979; PubMed Central PMCID: PMC11571630.

9. Drucker DJ. GLP-1-based therapies for diabetes, obesity and beyond. Nat Rev Drug Discov. 2025 Aug;24(8):631–50. doi:10.1038/s41573-025-01183-8

10. Clifton PG. Meal patterning in rodents: psychopharmacological and neuroanatomical studies. Neurosci Biobehav Rev. 2000 Mar 1;24(2):213–22. doi:10.1016/S0149-7634(99)00074-3

11. Ali MA, Kravitz AV. Challenges in Quantifying Food Intake in Rodents. Brain Res. 2018 Aug 15;1693(Pt B):188–91. doi:10.1016/j.brainres.2018.02.040 PubMed PMID: 29903621; PubMed Central PMCID: PMC6005200.

12. Shao G, Helu M. Framework for a digital twin in manufacturing: Scope and requirements. Manuf Lett. 2020 Apr 1;24:105–7. doi:10.1016/j.mfglet.2020.04.004

13. Katsoulakis E, Wang Q, Wu H, Shahriyari L, Fletcher R, Liu J, et al. Digital twins for health: a scoping review. Npj Digit Med. 2024 Mar 22;7(1):1–11. doi:10.1038/s41746-024-01073-0

14. Sharma A, Kosasih E, Zhang J, Brintrup A, Calinescu A. Digital Twins: State of the art theory and practice, challenges, and open research questions. J Ind Inf Integr. 2022 Nov 1;30:100383. doi:10.1016/j.jii.2022.100383

15. Knight-Schrijver VR, Chelliah V, Cucurull-Sanchez L, Le Novère N. The promises of quantitative systems pharmacology modelling for drug development. Comput Struct Biotechnol J. 2016 Jan 1;14:363–70. doi:10.1016/j.csbj.2016.09.002

16. Bradshaw EL, Spilker ME, Zang R, Bansal L, He H, Jones RDO, et al. Applications of Quantitative Systems Pharmacology in Model-Informed Drug Discovery: Perspective on Impact and Opportunities. CPT Pharmacomet Syst Pharmacol. 2019 Nov;8(11):777–91. doi:10.1002/psp4.12463 PubMed PMID: 31535440; PubMed Central PMCID: PMC6875708.

17. Gennemark P, Trägårdh M, Lindén D, Ploj K, Johansson A, Turnbull A, et al. Translational Modeling to Guide Study Design and Dose Choice in Obesity Exemplified by AZD1979, a Melanin-concentrating Hormone Receptor 1 Antagonist. CPT Pharmacomet Syst Pharmacol. 2017 Jul;6(7):458–68. doi:10.1002/psp4.12199 PubMed PMID: 28556607; PubMed Central PMCID: PMC5529746.

18. Brown A, Tzanakakis ES. Mathematical modeling clarifies the paracrine roles of insulin and glucagon on the glucose-stimulated hormonal secretion of pancreatic alpha- and beta-cells. Front Endocrinol. 2023;14:1212749. doi:10.3389/fendo.2023.1212749 PubMed PMID: 37645413; PubMed Central PMCID: PMC10461634.

19. Palumbo P, Ditlevsen S, Bertuzzi A, De Gaetano A. Mathematical modeling of the glucose-insulin system: a review. Math Biosci. 2013 Aug;244(2):69–81. doi:10.1016/j.mbs.2013.05.006 PubMed PMID: 23733079.

20. De Gaetano A, Hardy T, Beck B, Abu-Raddad E, Palumbo P, Bue-Valleskey J, et al. Mathematical models of diabetes progression. Am J Physiol Endocrinol Metab. 2008 Dec;295(6):E1462–1479. doi:10.1152/ajpendo.90444.2008 PubMed PMID: 18780774.

21. Guo J, Hall KD. Predicting Changes of Body Weight, Body Fat, Energy Expenditure and Metabolic Fuel Selection in C57BL/6 Mice. PLOS ONE. 2011 Jan 5;6(1):e15961. doi:10.1371/journal.pone.0015961

22. Gennemark P, Jansson-Löfmark R, Hyberg G, Wigstrand M, Kakol-Palm D, Håkansson P, et al. A modeling approach for compounds affecting body composition. J Pharmacokinet Pharmacodyn. 2013 Dec;40(6):651–67. doi:10.1007/s10928-013-9337-x PubMed PMID: 24158456.

23. Herrgårdh T, Li H, Nyman E, Cedersund G. An Updated Organ-Based Multi-Level Model for Glucose Homeostasis: Organ Distributions, Timing, and Impact of Blood Flow. Front Physiol. 2021 Jun 1;12. doi:10.3389/fphys.2021.619254

24. Simonsson C, Lövfors W, Bergqvist N, Nyman E, Gennemark P, Stenkula KG, et al. A multi-scale *in silico* mouse model for diet-induced insulin resistance. Biochem Eng J. 2023 Feb 1;191:108798. doi:10.1016/j.bej.2022.108798

25. Silfvergren O, Rigal S, Schimek K, Simonsson C, Kanebratt KP, Forschler F, et al. M4 drug discovery: Human drug predictions from integrated preclinical insights exemplified with a GLP-1R agonist. Eur J Pharm Sci. 2026 Aug 30;107648. doi:10.1016/j.ejps.2026.107648

26. Kutumova E, Kiselev I, Sharipov R, Lifshits G, Kolpakov F. Thoroughly Calibrated Modular Agent-Based Model of the Human Cardiovascular and Renal Systems for Blood Pressure Regulation in Health and Disease. Front Physiol. 2021 Nov 11;12. doi:10.3389/fphys.2021.746300

27. European Union. Directive 2010/63/EU of the European Parliament and of the Council of 22 September 2010 on the protection of animals used for scientific purposes [Internet]. Official Journal of the European Union: European Union; 2010 Sep. Report 2010/63/EU. Available from: https://eur-lex.europa.eu/eli/dir/2010/63/oj/eng

28. Mack CM, Moore CX, Jodka CM, Bhavsar S, Wilson JK, Hoyt JA, et al. Antiobesity action of peripheral exenatide (exendin-4) in rodents: effects on food intake, body weight, metabolic status and side-effect measures. Int J Obes. 2006 Sep;30(9):1332–40. doi:10.1038/sj.ijo.0803284

29. He MQ, Wang JY, Wang Y, Sui J, Zhang M, Ding X, et al. High-fat diet-induced adipose tissue expansion occurs prior to insulin resistance in C57BL/6J mice. Chronic Dis Transl Med. 2020 Sep;6(3):198–207. doi:10.1016/j.cdtm.2020.06.003 PubMed PMID: 32885155; PubMed Central PMCID: PMC7451745.

30. Liang R, Feng X, Shi D, Yang M, Yu L, Liu W, et al. The global burden of disease attributable to high fasting plasma glucose in 204 countries and territories, 1990–2019: An updated analysis for the Global Burden of Disease Study 2019. Diabetes Metab Res Rev. 2022;38(8):e3572. doi:10.1002/dmrr.3572

31. American Diabetes Association Professional Practice Committee for Diabetes*. 2. Diagnosis and Classification of Diabetes: Standards of Care in Diabetes—2026. Diabetes Care. 2025 Dec 8;49(Supplement_1):S27–49. doi:10.2337/dc26-S002

32. Huang X, Zheng Q. Pharmacokinetic and Pharmacodynamic Data Analysis: Concepts and Applications. Am J Pharm Educ. 2010 Apr 12;74(3):53b. PubMed PMID: null; PubMed Central PMCID: PMC2865421.

33. Mager DE, Jusko WJ. General Pharmacokinetic Model for Drugs Exhibiting Target-Mediated Drug Disposition. J Pharmacokinet Pharmacodyn. 2001 Dec 1;28(6):507–32. doi:10.1023/A:1014414520282

34. Iwasaki S, Hamada T, Chisaki I, Andou T, Sano N, Furuta A, et al. Mechanism-Based Pharmacokinetic/Pharmacodynamic Modeling of the Glucagon-Like Peptide-1 Receptor Agonist Exenatide to Characterize Its Antiobesity Effects in Diet-Induced Obese Mice. J Pharmacol Exp Ther. 2017 Sep;362(3):441–9. doi:10.1124/jpet.117.242651 PubMed PMID: 28698254.

35. Thorens B, Porret A, Bühler L, Deng SP, Morel P, Widmann C. Cloning and functional expression of the human islet GLP-1 receptor. Demonstration that exendin-4 is an agonist and exendin-(9-39) an antagonist of the receptor. Diabetes. 1993 Nov;42(11):1678–82. doi:10.2337/diab.42.11.1678 PubMed PMID: 8405712.

36. Wild D, Béhé M, Wicki A, Storch D, Waser B, Gotthardt M, et al. [Lys40(Ahx-DTPA-111In)NH2]exendin-4, a very promising ligand for glucagon-like peptide-1 (GLP-1) receptor targeting. J Nucl Med Off Publ Soc Nucl Med. 2006 Dec;47(12):2025–33. PubMed PMID: 17138746.

37. Almoshari Y. Osmotic Pump Drug Delivery Systems—A Comprehensive Review. Pharmaceuticals. 2022 Nov;15(11):1430. doi:10.3390/ph15111430

38. McLane VD, Bergquist I, Cormier J, Barlow DJ, Houseknecht KL, Bilsky EJ, et al. Long-Term Morphine Delivery via Slow Release Morphine Pellets or Osmotic Pumps: Plasma Concentration, Analgesia, and Naloxone-Precipitated Withdrawal. Life Sci. 2017 Sep 15;185:1–7. doi:10.1016/j.lfs.2017.07.016 PubMed PMID: 28723417; PubMed Central PMCID: PMC5577921.

39. Shah H, Ayala JE. Prolonged Semaglutide Treatment Reveals Stage-Dependent Changes to Feeding Behavior and Metabolic Adaptations in Male Mice. Diabetes. 2025 Dec 2;75(2):288–300. doi:10.2337/db25-0678

40. Hsu TM, Hahn JD, Konanur VR, Lam A, Kanoski SE. Hippocampal GLP-1 Receptors Influence Food Intake, Meal Size, and Effort-Based Responding for Food through Volume Transmission. Neuropsychopharmacology. 2015 Jan;40(2):327–37. doi:10.1038/npp.2014.175 PubMed PMID: 25035078; PubMed Central PMCID: PMC4443945.

41. Gong Z, Chen R, Ye L, Ma G, Xian Y, Kulkarni K. Editorial: Pharmacokinetic-pharmacodynamic model of drugs and their pharmacokinetic differences between normal and disease states. Front Pharmacol. 2024 Aug 5;15:1451176. doi:10.3389/fphar.2024.1451176 PubMed PMID: 39161901; PubMed Central PMCID: PMC11330815.

42. Moretto TL, Benfato ID, de Carvalho FP, Barthichoto M, Le Sueur-Maluf L, de Oliveira CAM. The effects of calorie-matched high-fat diet consumption on spontaneous physical activity and development of obesity. Life Sci. 2017 Jun 15;179:30–6. doi:10.1016/j.lfs.2017.04.017 PubMed PMID: 28449870.

43. Bjursell M, Gerdin AK, Lelliott CJ, Egecioglu E, Elmgren A, Törnell J, et al. Acutely reduced locomotor activity is a major contributor to Western diet-induced obesity in mice. Am J Physiol Endocrinol Metab. 2008 Feb;294(2):E251–260. doi:10.1152/ajpendo.00401.2007 PubMed PMID: 18029443.

44. Maric I, Krieger JP, van der Velden P, Börchers S, Asker M, Vujicic M, et al. Sex and Species Differences in the Development of Diet-Induced Obesity and Metabolic Disturbances in Rodents. Front Nutr. 2022;9:828522. doi:10.3389/fnut.2022.828522 PubMed PMID: 35284452; PubMed Central PMCID: PMC8908889.

45. Melhorn SJ, Krause EG, Scott KA, Mooney MR, Johnson JD, Woods SC, et al. Acute exposure to a high-fat diet alters meal patterns and body composition. Physiol Behav. 2010 Jan 12;99(1):33–9. doi:10.1016/j.physbeh.2009.10.004 PubMed PMID: 19835896; PubMed Central PMCID: PMC2794977.

46. Atalayer D, Rowland NE. Meal patterns of mice under systematically varying approach and unit costs for food in a closed economy. Physiol Behav. 2009 Aug 4;98(1–2):85–93. doi:10.1016/j.physbeh.2009.04.016 PubMed PMID: 19394352; PubMed Central PMCID: PMC2722741.

47. Rathod YD, Di Fulvio M. The feeding microstructure of male and female mice. PLoS ONE. 2021 Feb 4;16(2):e0246569. doi:10.1371/journal.pone.0246569 PubMed PMID: 33539467; PubMed Central PMCID: PMC7861458.

48. Ren B, Huang Y, Zhang J, Li J, Liu Z, Guan Y, et al. Impact of Time-Restricted Feeding on Adaptation to a 6-Hour Delay Phase Shift or a 12-Hour Phase Shift in Mice. Nutrients. 2022 Jul 23;14(15):3025. doi:10.3390/nu14153025 PubMed PMID: 35893879; PubMed Central PMCID: PMC9329972.

49. Fonken LK, Workman JL, Walton JC, Weil ZM, Morris JS, Haim A, et al. Light at night increases body mass by shifting the time of food intake. Proc Natl Acad Sci U S A. 2010 Oct 26;107(43):18664–9. doi:10.1073/pnas.1008734107 PubMed PMID: 20937863; PubMed Central PMCID: PMC2972983.

50. Dalla Man C, Rizza RA, Cobelli C. Meal simulation model of the glucose-insulin system. IEEE Trans Biomed Eng. 2007 Oct;54(10):1740–9. doi:10.1109/TBME.2007.893506 PubMed PMID: 17926672.

51. Stahel P, Cant JP, MacPherson JAR, Berends H, Steele MA. A Mechanistic Model of Intermittent Gastric Emptying and Glucose-Insulin Dynamics following a Meal Containing Milk Components. PLoS ONE. 2016 Jun 2;11(6):e0156443. doi:10.1371/journal.pone.0156443 PubMed PMID: 27253712; PubMed Central PMCID: PMC4890795.

52. Yokrattanasak J, Gaetano AD, Panunzi S, Satiracoo P, Lawton WM, Lenbury Y. A Simple, Realistic Stochastic Model of Gastric Emptying. PLOS ONE. 2016 Apr 8;11(4):e0153297. doi:10.1371/journal.pone.0153297

53. Jacome-Sosa M, Hu Q, Manrique-Acevedo CM, Phair RD, Parks EJ. Human intestinal lipid storage through sequential meals reveals faster dinner appearance is associated with hyperlipidemia. JCI Insight. 2021 Aug 9;6(15):e148378. doi:10.1172/jci.insight.148378 PubMed PMID: 34369385; PubMed Central PMCID: PMC8489663.

54. Silfvergren O, Simonsson C, Ekstedt M, Lundberg P, Gennemark P, Cedersund G. Digital twin predicting diet response before and after long-term fasting. PLOS Comput Biol. 2022 Sep 12;18(9):e1010469. doi:10.1371/journal.pcbi.1010469

55. Simonsson C, Silfvergren O, Podéus H, Tunedal K, Lövfors W, Stenkula KG, et al. A mathematical model of dietary lipid absorption and postprandial chylomicron dynamics. Metab Open. 2026 Dec 1;32:100522. doi:10.1016/j.metop.2026.100522

56. Jansson-Löfmark R, Hjorth S, Gabrielsson J. Does In Vitro Potency Predict Clinically Efficacious Concentrations? Clin Pharmacol Ther. 2020 Aug;108(2):298–305. doi:10.1002/cpt.1846 PubMed PMID: 32275768; PubMed Central PMCID: PMC7484912.

57. Boianelli A, Nordell P, Earl J, Naylor J, Hornigold D, Jansson Löfmark R, et al. Establishing a Relationship between In Vitro Potency in Cell-Based Assays and Clinical Efficacious Concentrations for Approved GLP-1 Receptor Agonists. Pharmaceutics. 2024 Oct 8;16(10):1310. doi:10.3390/pharmaceutics16101310 PubMed PMID: 39458639; PubMed Central PMCID: PMC11510446.

58. Brännmark C, Nyman E, Fagerholm S, Bergenholm L, Ekstrand EM, Cedersund G, et al. Insulin signaling in type 2 diabetes: experimental and modeling analyses reveal mechanisms of insulin resistance in human adipocytes. J Biol Chem. 2013 Apr 5;288(14):9867–80. doi:10.1074/jbc.M112.432062 PubMed PMID: 23400783; PubMed Central PMCID: PMC3617287.

59. Nyman E, Brännmark C, Palmér R, Brugård J, Nyström FH, Strålfors P, et al. A hierarchical whole-body modeling approach elucidates the link between in Vitro insulin signaling and in Vivo glucose homeostasis. J Biol Chem. 2011 Jul 22;286(29):26028–41. doi:10.1074/jbc.M110.188987 PubMed PMID: 21572040; PubMed Central PMCID: PMC3138269.

60. Cobelli C, Dalla Man C, Toffolo G, Basu R, Vella A, Rizza R. The oral minimal model method. Diabetes. 2014 Apr;63(4):1203–13. doi:10.2337/db13-1198 PubMed PMID: 24651807; PubMed Central PMCID: PMC4179313.

61. Chudtong M, Gaetano AD. A mathematical model of food intake. Math Biosci Eng MBE. 2021 Jan 15;18(2):1238–79. doi:10.3934/mbe.2021067 PubMed PMID: 33757185.

62. Chudtong M, Gaetano AD. Food choices and body weight changes: A mathematical model analysis. Math Biosci Eng MBE. 2025 Jun 5;22(7):1790–824. doi:10.3934/mbe.2025065 PubMed PMID: 40676993.

63. Cedersund G. Conclusions via unique predictions obtained despite unidentifiability--new definitions and a general method. FEBS J. 2012 Sep;279(18):3513–27. doi:10.1111/j.1742-4658.2012.08725.x PubMed PMID: 22846178.

64. Cedersund G, Roll J. Systems biology: model based evaluation and comparison of potential explanations for given biological data. FEBS J. 2009;276(4):903–22. doi:10.1111/j.1742-4658.2008.06845.x

65. Podéus H, Magnusson G, Keshmiri S, Tunedal K, Sundqvist N, Lövfors W, et al. SUND: simulation using nonlinear dynamic models - a toolbox for simulating multi-level, time-dynamic systems in a modular way [Internet]. arXiv; 2025 [cited 2026 Sep 11]. Available from: https://arxiv.org/abs/2510.13932 doi:10.48550/ARXIV.2510.13932

66. Silfvergren O, Rigal S, Schimek K, Simonsson C, Kanebratt KP, Forschler F, et al. An Integrated Analysis of GLP-1R Agonist Mechanisms: Addressing Study Variations in Heterogeneous Cell Systems [Internet]. bioRxiv; 2026 [cited 2026 Apr 2]. p. 2026.03.25.714139. Available from: https://www.biorxiv.org/content/10.64898/2026.03.25.714139v1 doi:10.64898/2026.03.25.714139

67. Annabel L. M, Bernice K. W. Energy Value of Foods: Basis and Derivation. Washington, DC: United States Department of Agriculture; 1955. Report Agriculture Handbook No. 74.

68. Whited KL, Hornof WJ, Garcia T, Bohan DC, Larson RF, Raybould HE. A non-invasive method for measurement of gastric emptying in mice: effects of altering fat content and CCK A receptor blockade. Neurogastroenterol Motil. 2004 Aug 1;16(4):421–7. doi:10.1111/j.1365-2982.2004.00529.x PubMed PMID: 15305997.

69. Symonds E, Butler R, Omari T. Noninvasive breath tests can detect alterations in gastric emptying in the mouse. Eur J Clin Invest. 2002 May;32(5):341–4. doi:10.1046/j.1365-2362.2002.00991.x PubMed PMID: 12027874.

70. Miller KE, Bajzer Ž, Hein SS, Phillips JE, Syed S, Wright AM, et al. High temporal resolution gastric emptying breath tests in mice. Neurogastroenterol Motil. 2018 Mar 1;e13333. doi:10.1111/nmo.13333 PubMed PMID: 29575442; PubMed Central PMCID: PMC6157017.

71. WebPlotDigitizer - Copyright 2010-2024 Ankit Rohatgi [Internet]. [cited 2025 Apr 8]. Available from: https://apps.automeris.io/wpd4/

72. Gao W, Jusko WJ. Target-mediated pharmacokinetic and pharmacodynamic model of exendin-4 in rats, monkeys, and humans. Drug Metab Dispos Biol Fate Chem. 2012 May;40(5):990–7. doi:10.1124/dmd.111.042291 PubMed PMID: 22338110; PubMed Central PMCID: PMC3336795.

