## Supplementary Note S1 for "A Mathematical Model Connecting Appetite-Driven Food Intake to Metabolic Changes across Diet and GLP-1RA Treatments in Mice"

We describe the effects of diet and GLP-1RAs on eating behaviours and their downstream consequences by connecting a new appetite model to pre-existing body mass (1,2), glucose-insulin (3), and GLP-1RA (4) models. Minimal changes were made to these models to incorporate GLP-1RA effects, enable interactions among the modules, and allow their application to the new scenarios considered in this study.

In this supplementary note these changes are described one-by-one, together with a detailed description of each model, beginning with the glucose-insulin module.

### The glucose-insulin module

The glucose-insulin module is a further development of a pre-existing model describing glucose responses to meals without GLP-1RA effects (3). To incorporate GLP-1RA effects, equations from our previously developed model were imported (4). The key changes made include: the digestive system was replaced by the appetite module; insulin secretion was updated, which now has a glucose-dependent and -independent part; and an effect of GLP-1RAs on glucose-dependent insulin secretion was added.

The module consists of six ordinary differential equations (ODEs) describing three glucose states, one insulin state, and two delayed regulatory states. All model-state derivatives were multiplied by 24 to convert the time unit of the original model from hours to days, which is the time unit used in all models. The stated initial values were used for a steady-state simulation. All model states were simulated to steady state before the experimental setting-specific simulations.

The first ODE describes the amount of glucose in the subcutaneous compartment:

$$\frac{d}{dt}(G_{SC}) = -G_{SC} * k_{ip} \qquad \text{(Equation S1)}$$

$$G_{SC}\,(0) = 0 \text{ mmol} \qquad \text{(Equation S2)}$$

where $k_{ip}$ is the transfer rate of glucose from the subcutaneous compartment to the central glucose compartment.

The second glucose state describes the amount of glucose in the central compartment:

$$\frac{d}{dt}(G_{Central}) = G_{SC} * k_{ip} + Meal_{glucose} + QVP * G_{Peripheral} - QVG * G_{central} - U + EGP \qquad \text{(Equation S3)}$$

$$G_{Central}\,(0) = 0.0179 \text{ mmol} \qquad \text{(Equation S4)}$$

where $G_{SC} * k_{ip}$ is the appearance of subcutaneously administered glucose; $Meal_{glucose}$ is glucose from meals provided by the appetite module explained in the main text (**Eq. 15**); QVP is a model function representing glucose transfer rate between the peripheral and central compartment; $G_{Peripheral}$ is a model state representing glucose in the peripheral compartment; U is a model function representing glucose utilisation; and EGP is a model function representing endogenous glucose production.

Plasma glucose concentration is calculated from the central glucose amount as:

$$Plasma_{glucose} = \frac{G_{central}}{VG} \qquad \text{(Equation S5)}$$

where VG is the central glucose distribution volume. In accordance with the original model, VG was set to 0.003065 L. Together with the initial central glucose amount before the steady-state simulation, this gives a plasma glucose concentration of approximately 5.84 mmol/L.

The third glucose state describes the amount of glucose in the peripheral compartment:

$$\frac{d}{dt}\left(G_{Peripheral}\right) = QVG * G_{central} - QVP * G_{Peripheral} \qquad \text{(Equation S6)}$$

$$G_{Peripheral}\ (0) = 0.0165\ \text{mmol} \qquad \text{(Equation S7)}$$

where $QVG * G_{central} - QVP * G_{Peripheral}$ represents the glucose transfer rate between the peripheral and central compartments introduced in the previously listed ODE for $G_{Central}$ (**Eq. S3**).

The compartment-specific fractional transport rates are:

$$QVP = \frac{Q}{VP} \qquad \text{(Equation S8)}$$

$$QVG = \frac{Q}{VG} \qquad \text{(Equation S9)}$$

where Q is a model parameter representing intercompartmental glucose flow; VP is the peripheral glucose distribution volume; and VG is the central glucose distribution volume. The same distribution volumes as in the original model were used ($VP = 0.00281$ L; $VG = 0.003065$ L)

The fourth ODE describes the amount of insulin in the central insulin compartment:

$$\frac{d}{dt}\left(I_{Central}\right) = IP - CL_I * Plasma_{insulin} \qquad \text{(Equation S10)}$$

$$I_{Central}\ (0) = 0.13766\ \text{pmol} \qquad \text{(Equation S11)}$$

where IP is a model function representing insulin production rate; $CL_I$ is a model parameter representing insulin clearance; and $Plasma_{insulin}$ is insulin concentration in the central compartment, which was calculated as:

$$Plasma_{insulin} = \frac{I_{central}}{VI} \qquad \text{(Equation S12)}$$

where VI is the central insulin distribution volume set to 0.002001 L, in accordance with the original model. The initial values of $I_{central}$ and VI provide an initial insulin concentration of 68.8 pmol/L.

Insulin production is described as:

$$IP = (I_{SS} + I_{Gdep} * Exenatide_{PD} * \max(0, Plasma_{glucose} - I_{G0} * fIR_{EGP})) * fIR_{Ins} \qquad \text{(Equation S13)}$$

Where $I_{SS}$ is a model parameter representing the basal insulin production rate; $I_{Gdep}$ is a model parameter representing glucose-stimulated insulin production; $Exenatide_{PD}$ is a model function describing the effect of exenatide on glucose-stimulated insulin secretion; $I_{G0}$ is a model parameter representing the plasma glucose threshold above which glucose-dependent insulin production occurs; $fIR_{EGP}$ is a model function that modifies the glucose threshold in response to changes in basal glucose caused by metabolic dysregulation; and $fIR_{Ins}$ is a model function that modifies total insulin production in response to metabolic dysregulation. The functions $fIR_{EGP}$ and $fIR_{Ins}$ are inputs supplied by the body mass module and are described in the subsequent section on that module.

As previously mentioned, the model function IP was modified using components from the previous model with GLP-1RA support (4). The separation of basal and stimulated insulin secretion is new and was introduced to allow exenatide to affect only glucose-stimulated insulin secretion through the term $Exenatide_{PD}$. The $fIR_{EGP}$ term is from the original glucose-insulin model, and was added to account for changes in basal plasma glucose concentration.

The fifth ODE describes a delayed insulin concentration used to regulate insulin-dependent glucose utilisation:

$$\frac{d}{dt}(U_{IdepDelay}) = K_{IE} * (Plasma_{insulin} - U_{IdepDelay}) \quad \text{(Equation S14)}$$

$$U_{IdepDelay}(0) = 68.8 \text{ pmol/L} \quad \text{(Equation S15)}$$

Where $K_{IE}$ is a model parameter determining the delay between plasma insulin concentration and insulin-dependent glucose utilisation.

The sixth and final ODE describes a delayed glucose concentration used to regulate endogenous glucose production:

$$\frac{d}{dt}(EGP_{GdepDelay}) = K_{GE2} * (Plasma_{glucose} - EGP_{GdepDelay}) \quad \text{(Equation S16)}$$

$$EGP_{GdepDelay}(0) = 5.85 \text{ mmol/L} \quad \text{(Equation S17)}$$

where $K_{GE2}$ is a model parameter determining the delay between plasma glucose concentration and endogenous glucose production.

Several ODEs described above depend on model functions. The total glucose utilisation in Equation S3 is described as the sum of glucose utilisation through different pathways:

$$U = CL_G \cdot Plasma_{glucose} + Uid \quad \text{(Equation S18)}$$

where $CL_G$ is a model parameter representing insulin-independent glucose clearance by, for example, the brain, and Uid is a model function representing insulin-dependent glucose utilisation:

$$Uid = U_{Max} \cdot \frac{Plasma_{glucose}}{U_{50} + Plasma_{glucose}} \quad \text{(Equation S19)}$$

$$U_{Max} = U_{Max0} + U_{Maxid} * \frac{U_{IdepDelay}}{fIR_{Uid}} \quad \text{(Equation S20)}$$

where $U_{50}$ is a model parameter representing the plasma glucose concentration at which insulin-dependent glucose utilisation reaches half of maximum utilization termed $U_{Max}$; $U_{Max0}$ is a model parameter representing basal insulin-dependent glucose utilisation; $U_{Maxid}$ is a model parameter that increases glucose utilisation in response to the delayed insulin signal $U_{IdepDelay}$ (**Eq. S14**); and $fIR_{Uid}$is a model input from the body mass module that describes insulin resistance. In the previous works (5–7), Uid was separated into organ-specific utilisation using organ-specific equations. In the present model, it was instead represented by fixed ratios to minimise changes to the original model and because the model was not fitted to organ-specific glucose-utilisation data: adipose tissue (10%), skeletal muscle (40%), and liver (50%).

The endogenous glucose production is described as:

$$EGP = \frac{G_{ss} * fIR_{EGP}}{\left(\frac{EGP_{GdepDelay}}{G_{ss} * fIR_{EGP}}\right)^{GPRG}} \quad \text{(Equation S21)}$$

where $G_{ss}$ is the model parameter governing endogenous glucose production in the absence of metabolic dysregulation; $fIR_{EGP}$ is a model function that increases $G_{ss}$ in response to metabolic dysregulation; $EGP_{GdepDelay}$ is the delayed plasma glucose concentration (Eq. S16), which drives glucose concentrations towards baseline by reducing production at higher glucose concentrations; and GPRG is a model parameter determining the non-linearity of glucose-dependent suppression of endogenous glucose production.

The pharmacodynamic effect of GLP-1RA on stimulated insulin secretion is calculated as:

$$GLP1RA_{PD} = 1 + \frac{GLP1RA_{Max} * Plasma_{GLP1RA}^{GLP1RA_{Hill}}}{GLP1RA_{50}^{GLP1RA_{Hill}} + Plasma_{GLP1RA}^{GLP1RA_{Hill}}} - \frac{GLP1RA\_Decay_{Max} * Plasma_{GLP1RA}^{GLP1RA_{Hill}}}{GLP1RA\_Decay_{50}^{GLP1RA_{Hill}} + Plasma_{GLP1RA}^{GLP1RA_{Hill}}} \quad \text{(Equation S22)}$$

where $GLP1RA_{PD}$ describes the bell-shaped GLP-1RA exposure-effect relationship on glucose-stimulated insulin secretion; $GLP1RA_{Max}$ and $GLP1RA\_Decay_{Max}$ are model parameters that represent the maximum effect of both the positive and negative parts of the bell-shaped dose-response curve; $GLP1RA_{Hill}$ is a model parameter representing the Hill coefficient from no response to maximum response; $GLP1RA_{50}$ and $GLP1RA\_Decay_{50}$ are model parameters describing the $EC_{50}$ and $IC_{50}$ of the bell-shaped exposure-effect relationship; and $Plasma_{GLP1RA}$ is the concentration of GLP-1RA in the central compartment.

The glucose-insulin module uses five external inputs:

$$Plasma_{GLP1RA} = C1(t) \quad \text{(Equation S23)}$$
$$fIR_{Uid} = fIR_{Uid}(t) \quad \text{(Equation S24)}$$
$$fIR_{EGP} = fIR_{EGP}(t) \quad \text{(Equation S25)}$$
$$fIR_{Ins} = fIR_{Ins}(t) \quad \text{(Equation S26)}$$
$$Meal_{glucose} = Meal_{glucose}(t) \quad \text{(Equation S27)}$$

where C1 is the plasma GLP-1RA concentration supplied by the GLP-1RA module; $fIR_{Uid}$, $fIR_{EGP}$, and $fIR_{Ins}$ are the three functions describing changes in metabolic dysregulation, supplied by the body mass module; and $Meal_{glucose}$ is the glucose appearance rate supplied by the appetite module. In the absence of external signals, $Plasma_{GLP1RA}$ and $Meal_{glucose}$ are set to zero, while the three metabolic-dysregulation inputs are set to one.

The parameters used in this module are presented below.

| **Parameter** | **Fitted value** best (min-max) | **Unit** |
|---|---|---|
| **Q** | 0.02 (0.01–0.05) | L/h |
| $\mathbf{K_{IE}}$ | 0.17 (0.06–0.25) | 1/h |
| $\mathbf{K_{GE2}}$ | 23.24 (23.24–46.80) | 1/h |
| $\mathbf{CL_{G}}$ | 0.03 (0.01–0.03) | L/h |
| $\mathbf{k_{ip}}$ | 1.64 (1.24–1.85) | 1/h |
| **GPRG** | 2.00 (1.68–2.00) | dimensionless |
| $\mathbf{U_{50}}$ | 20.00 (19.24–20.00) | mmol/L |
| $\mathbf{U_{Maxid}}$ | 0.0067 (0.0067–0.0097) | mmol/(h·pmol/L) |
| $\mathbf{U_{Max0}}$ | 0.77 (0.22–0.87) | mmol/h |
| $\mathbf{G_{ss}}$ | 4.62 (3.77–5.16) | mmol/h |
| $\mathbf{I_{SS}}$ | 63.68 (52.70–63.68) | pmol/h |
| $\mathbf{I_{Gdep}}$ | 5.70 (4.74–5.70) | pmol/(h·mmol/L) |
| $\mathbf{I_{G0}}$ | 11.83 (10.29–14.21) | mmol/L |
| $\mathbf{CL_{I}}$ | 0.22 (0.19–0.22) | L/h |
| $\mathbf{GLP1RA_{Max}}$ | 8.68 (6.16–10.00) | fold-increase |
| $\mathbf{GLP1RA_{50}}$ | 1.35 (0.71–1.94) | nmol/L |
| $\mathbf{GLP1RA_{Hill}}$ | 4.00 (2.00–4.00) | dimensionless |
| $\mathbf{GLP1RA\_Decay_{Max}}$ | 7.25 (4.47–7.25) | fold-decrease |
| $\mathbf{GLP1RA\_Decay_{50}}$ | 7.60 (4.71–10.97) | nmol/L |

### The GLP-1RA module

The GLP-1RA module describes the pharmacokinetics of exenatide and semaglutide and is based on our previously developed model (4). The module consists of three ordinary differential equations: one state describing the amount of GLP-1RA at the subcutaneous administration site and two states describing GLP-1RA concentrations in the central and peripheral compartments. The original rate equations were expressed per minute and were multiplied by 1440 to convert the model time unit to days, which is the time unit used in all modules. All states were initialised to zero before each treatment simulation.

The first ODE describes the amount of GLP-1RA at the subcutaneous administration site:

$$\frac{d}{dt}(A) = SC_{infusion} - \frac{Ka * A}{Ka50 + A} \quad \text{(Equation S28)}$$

$$A(0) = 0 \text{ nmol} \quad \text{(Equation S29)}$$

where $SC_{infusion}$ is a model input representing subcutaneous infusion; Ka is a model parameter describing the maximum absorption rate from the subcutaneous administration site to the central compartment; and Ka50 is the amount at the subcutaneous administration site at which absorption reaches half its maximum rate.

The second ODE describes the GLP-1RA concentration in the central compartment:

$$\frac{d}{dt}(C1) = \frac{IV_{infusion}}{V1} + k21 * C2 * \frac{V2}{V1} + \frac{\frac{Ka * A}{Ka50 + A}}{V1} - \frac{\frac{CL_{Max} * C1}{CL_{50} + C1}}{V1} - k12 * C1 \quad \text{(Equation S30)}$$

$$C1(0) = 0 \text{ nmol/L} \quad \text{(Equation S31)}$$

where $IV_{infusion}$ is a model input representing intravenous infusion; V1 is a model function representing the central distribution volume; C2 is the concentration of GLP-1RA in the peripheral compartment; V2 is a model function representing the peripheral distribution volume; k12 and k21 are the intercompartmental fractional transport rates; $\frac{Ka*A}{Ka50+A}$ is absorption from the subcutaneous administration route; $CL_{Max}$ is a model parameter representing the maximum clearance; and $CL_{50}$ represents the concentration at which clearance reaches half its maximum rate.

The third ODE describes the GLP-1RA concentration in the peripheral compartment:

$$\frac{d}{dt}(C2) = k12 * C1 * \frac{V1}{V2} - k21 * C2 \quad \text{(Equation S32)}$$

$$C2(0) = 0 \text{ nmol/L} \quad \text{(Equation S33)}$$

where $k12 * C1 - k21 * C2$ represents transport between the central and peripheral compartments while accounting for differences in compartment volume $\frac{V1}{V2}$.

The central and peripheral distribution volumes are scaled to body mass in accordance with allometric scaling principles, sometimes referred to as 'Kleiber's law'(8):

$$V1 = V1_{ref} * \left(\frac{BW_{kg}}{BW_{ref}}\right)^{b_{volume}} \quad \text{(Equation S34)}$$

$$V2 = V2_{ref} * \left(\frac{BW_{kg}}{BW_{ref}}\right)^{b_{volume}} \quad \text{(Equation S35)}$$

where $V1_{ref}$ and $V2_{ref}$ are the reference central and peripheral distribution volumes; $BW_{kg}$ is the current body mass; $BW_{ref}$ is the reference body mass; and $b_{volume}$ is the model parameter describing the allometric exponent for the distribution volumes. The value of $b_{volume}$ was set to 1, in accordance with the pre-existing model.

Similarly, in accordance with the pre-existing PK model, clearance and intercompartmental flow are scaled to body mass:

$$CL = CL_{ref} * \left(\frac{BW_{kg}}{BW_{ref}}\right)^{b_{reaction}} \quad \text{(Equation S36)}$$

$$Q = Q_{ref} * \left(\frac{BW_{kg}}{BW_{ref}}\right)^{b_{reaction}} \quad \text{(Equation S37)}$$

where $CL_{ref}$ and $Q_{ref}$ are model parameters describing the maximum elimination rate and intercompartmental flow, respectively, and $b_{reaction}$ is the model parameter describing the allometric exponent for the reaction rates. The value of $b_{reaction}$ was set to 0.6, in accordance with the pre-existing model.

The compartment-specific fractional transport rates are calculated as:

$$k12 = \frac{Q}{V1} \quad \text{(Equation S38)}$$

$$k21 = \frac{Q}{V2} \quad \text{(Equation S39)}$$

where Q is a model parameter representing rate between the central and peripheral compartment.

The GLP-1RA module uses four external inputs:

$$BW_{kg} = BW_{kg}(t) \quad \text{(Equation S40)}$$

$$BW_{ref} = BW_{ref}(t) \quad \text{(Equation S41)}$$

$$IV_{Infusion} = IV_{Infusion}(t) \quad \text{(Equation S42)}$$

$$SC_{Infusion} = SC_{Infusion}(t) \quad \text{(Equation S43)}$$

where $BW_{kg}$ is the current body mass supplied by the body mass module or specified at the start of the simulation; $BW_{ref}$ is the reference body mass; and $IV_{Infusion}$ and $SC_{Infusion}$ are the intravenous and subcutaneous administration rates, respectively. In the absence of GLP-1RA treatment, both administration inputs are set to zero. $BW_{ref}$ was set to 0.34 kg when simulating exenatide, in accordance with the pre-existing model. $BW_{ref}$ was set to 0.05 kg when simulating semaglutide, representing the mice weight in the estimation data.

The GLP-1RA module provides one external output:

$$C1 = C1(t) \quad \text{(Equation S44)}$$

where C1 is the GLP-1RA concentration in the central compartment, which is supplied to the appetite and glucose-insulin modules to describe effects on appetite and glucose-stimulated insulin secretion, respectively.

The parameters used to describe exenatide in this module are presented below:

| Parameter | Fitted value<br>best (min-max) | Unit |
|---|---|---|
| **Ka** | 0.37 (0.07 - 1.74) | nmol/min |
| **Ka50** | 48.84 (8.83 - 220.77) | nmol |
| $\mathbf{Q_{ref}}$ | 0.0013 (0.0004 - 0.0034) | L/min |
| $\mathbf{V1_{ref}}$ | 0.06 (0.02 - 0.13) | L |
| $\mathbf{V2_{ref}}$ | 0.06 (0.03 - 0.11) | L |
| $\mathbf{CL_{ref}}$ | 2.24 (1.35 - 11.04) | nmol/min |
| $\mathbf{CL50_{ref}}$ | 309.82 (168.90 - 1503.97) | nmol/L |

To reflect the limited amount of pharmacokinetic data for semaglutide considered herein, several changes were made to simplify the model. For these simulations, $Q_{ref}$ was set to zero, which results in that the peripheral compartment is not used. Therefore, $V2_{ref}$ is not needed in the simulations and was not fitted to data. Furthermore, the non-linear clearance and absorption relationships used for the exenatide simulations were reduced to linear relationships for the semaglutide simulations. This was done by multiplying Ka, Ka50, $CL_{ref}$, and $CL50_{ref}$ by $10^9$ , and then optimising the ratios between Ka/Ka50, and $CL_{ref}/CL50_{ref}$, respectively. Multiplication by $10^9$ results in approximately linear relationships over the simulated ranges for the absorption rate (**Eq. S45**) and clearance (**Eq. S46**).

$$\text{Absorption}_{\text{semaglutide}} = \text{Ka} * \text{A} \quad \text{(Equation S45)}$$

$$\text{Clearance}_{\text{semaglutide}} = \text{CL}_{\text{ref}} * \frac{\text{C1}}{\text{V1}} \quad \text{(Equation S46)}$$

The choice to simplify the model for semaglutide reflects that these non-linear relationships could not be identified from the available estimation data and does not imply that maximum absorption or clearance rates do not exist.

The three parameters fitted to the semaglutide data are presented below:

| Parameter | Fitted value<br>best (min-max) | Unit |
|---|---|---|
| **Ka** | 24.91 (4.75 - 43.20) | 1/day |
| $\mathbf{V1_{ref}}$ | 0.012 (0.005 - 0.020) | L |
| $\mathbf{CL_{ref}}$ | 0.019 (0.015 - 0.026) | L/day |

### The body mass module

The body-mass module is based on previously published body-composition models describing changes in fat mass (FM), fat-free mass (FFM), and energy expenditure during body-weight perturbations. The body mass models were left unchanged (1,2), with an imported disease progression from the glucose-insulin model (3) and energy intake from the appetite module. The module consists of three ordinary differential equations: two states describing body composition, and one delayed regulatory state. The initial values for the body composition states were set to data, and the one delayed regulatory state was set using a steady state simulation. Time is expressed in days throughout the model.

The first ODE describes changes in fat-free mass:

$$\frac{d}{dt}(\text{FFM}) = \text{aa} * (\text{EI} - \text{EE}) \quad \text{(Equation S47)}$$

$$\text{FFM}(0) = 20.47 \text{ g} \quad \text{(Equation S48)}$$

where EI is a model input representing energy intake, which can be set to observation values or be predicted by the appetite module; EE is a model function representing energy expenditure, and aa is a model function determining the partitioning of energy imbalance into changes in fat-free mass.

The second ODE describes changes in fat mass:

$$\frac{d}{dt}(FM) = bb * (EI - EE) \quad \text{(Equation S49)}$$

$$FM(0) = 2.53\ g \quad \text{(Equation S50)}$$

where bb is a model function determining the partitioning of energy imbalance into changes in fat mass.

The final ODE describes a delayed adaptation signal:

$$\frac{d}{dt}(\lambda_{delay}) = \lambda_{delayK} * (\lambda - \lambda_{delay}) \quad \text{(Equation S51)}$$

$$\lambda_{delay}(0) = \lambda \quad \text{(Equation S52)}$$

where $\lambda$ is a model input representing energy expenditure from physical activity, and $\lambda_{delayK}$ is a model parameter governing the delay between $\lambda$ and its physiological effects.

Total body mass is calculated as:

$$BW = FM + FFM \quad \text{(Equation S53)}$$

and body mass in kilograms as:

$$BW_{kg} = \frac{BW}{1000} \quad \text{(Equation S54)}$$

The relative fat-mass and fat-free-mass fractions used by the insulin-resistance functions from the original glucose-insulin model (3) were imported:

$$xFM = \frac{FM}{FM_{init}} \quad \text{(Equation S55)}$$

$$xFFM = \frac{FFM}{FFM_{init}} \quad \text{(Equation S56)}$$

where $FM_{init}$ and $FFM_{init}$ are the initial values of fat mass and fat-free mass, respectively.

The energy-partitioning coefficient is calculated from:

$$\alpha = \alpha_c + \alpha_d \cdot \exp(\alpha_k \cdot FM) \quad \text{(Equation S57)}$$

where $\alpha_c$, $\alpha_d$, and $\alpha_k$ are model parameters from the original body mass models used to determine the partitioning of energy imbalance into changes in both fat-free mass and fat mass. In agreement with the subsequent expansion of the body mass model the values were allowed to be different depending on mouse sex (2).

The partitioning coefficients used in the body-composition equations are:

$$aa = \frac{\alpha}{\alpha * \rho_{FFM} + \rho_{FM}} \quad \text{(Equation S58)}$$

$$bb = \frac{1}{\alpha * \rho_{FFM} + \rho_{FM}} \quad \text{(Equation S59)}$$

where $\rho_{FFM}$ and $\rho_{FM}$ are the energy densities of fat mass and fat-free mass, respectively.

Energy expenditure is calculated from:

$$\Delta EI = EI - cEI \quad \text{(Equation S60)}$$

$$cc = K + \beta \cdot \Delta EI + (\gamma_{FFM} + \lambda_{delay}) \cdot FFM + (\gamma_{FM} + \lambda_{delay}) \cdot FM \quad \text{(Equation S61)}$$

$$EE = \frac{cc + \pi_{FM} \cdot bb \cdot EI + \pi_{FFM} \cdot aa \cdot EI}{1 + \pi_{FM} \cdot bb + \pi_{FFM} \cdot aa} \quad \text{(Equation S62)}$$

where cEI is a baseline energy intake, K is a baseline energy expenditure, β determines adaptive thermogenesis caused by changes in energy intake, $\gamma_{FFM}$and $\gamma_{FM}$ describe mass-dependent energy expenditure, and $\pi_{FM}$ and $\pi_{FFM}$ describe energetic costs associated with tissue turnover. As previously stated, no change was made to these equations and for a more in detailed description see the original publications.

A new addition to the body mass module is the three equations describing disease progression from changes in fat mass:

$$fIR_{Uid} = \frac{\log\left(1 + (DiseaseProgression_{scaling} * b_{CLGI})^3\ xFM\right)}{\log\left(1 + (DiseaseProgression_{scaling} * b_{CLGI})^3\right)} \quad \text{(Equation S63)}$$

$$fIR_{EGP} = \frac{\log\left(1 + (DiseaseProgression_{scaling} * b_{EGP})^3 \cdot xFM\right)}{\log\left(1 + (DiseaseProgression_{scaling} * b_{EGP})^3\right)} \quad \text{(Equation S64)}$$

$$fIR_{Ins} = \frac{\log\left(1 + (DiseaseProgression_{scaling} * b_{Ins})^3 \cdot xFM\right)}{\log\left(1 + (DiseaseProgression_{scaling} * b_{Ins})^3\right)} \quad \text{(Equation S65)}$$

Where $b_{CLGI}$, $b_{EGP}$, and $b_{Ins}$ are model parameters which determine the disease progression changes to xFM; 3 was used to increase parameter changes effect on the function at higher values; and $DiseaseProgression_{scaling}$ was one model parameter fitted to data. These three equations were revised from the original model, in which they were previously expressed using the general formula: $f(xFM) = 1 + parameter * \log(xFM)$. Where in the original model, $b_{CLGI}$, $b_{EGP}$, and $b_{Ins}$ were set to 0.4747, 0.1812, and 0.5391, respectively. At low positive values of xFM, this general formula can become negative, which provided unphysiological variables and crashes in our simulated interventions with large mass reductions. To avoid this, we revised the equations, so they always intercept both (x,y)=(1,1), in accordance to the previous general formula, and (x,y) = (0,0), to avoid crashing and unphysiological values.

To minimise difference between the original and revised functions $b_{CLGI}$, $b_{EGP}$, and $b_{Ins}$ were set to 4.8, 70, and 3.87, respectively. These three values were multiplied with the same model parameter termed $DiseaseProgression_{scaling}$ to account for changes between the original models and the revised multi-modular model. With these values, the revised functions provide only small differences to the original functions in the simulated intervals, with the important differences that y cannot become negative at low positive xFM values:

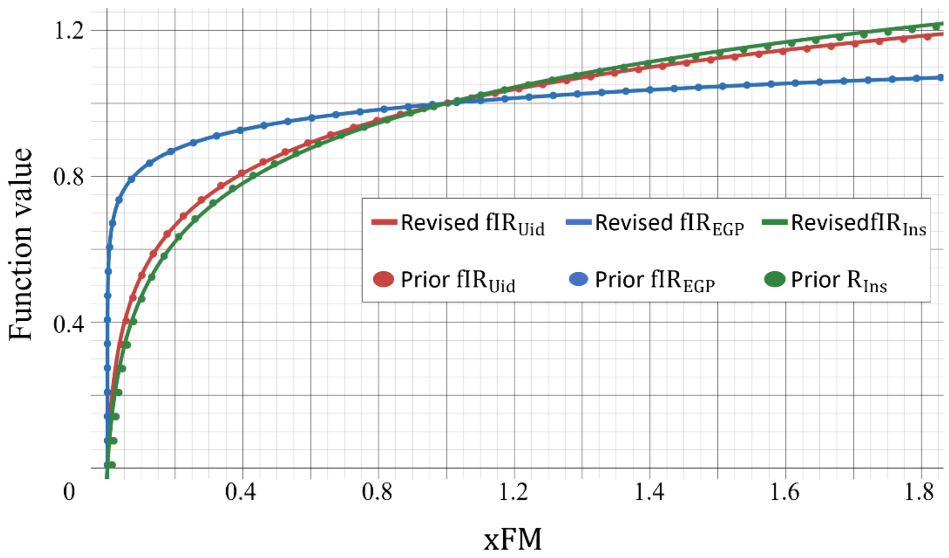


The body mass module uses five external inputs:

$$EI = EI(t) \quad \text{(Equation S66)}$$
$$sex = sex\,(t) \quad \text{(Equation S67)}$$
$$\lambda = \lambda(t) \quad \text{(Equation S68)}$$
$$FM_{init} = FM_{init}(t) \quad \text{(Equation S69)}$$
$$FFM_{init} = FFM_{init}(t) \quad \text{(Equation S70)}$$

where EI is energy intake provided by the appetite module, sex specifies the sex-specific parameterisation, λ represents energy expenditure associated with physical activity, and $FM_{init}$ and $FFM_{init}$ determine baseline body composition.

The body mass module uses four external outputs:

$$BW = BW(t) \quad \text{(Equation S71)}$$
$$fIR_{Uid} = fIR_{Uid}\,(t) \quad \text{(Equation S72)}$$
$$fIR_{EGP} = fIR_{EGP}(t) \quad \text{(Equation S73)}$$
$$fIR_{Ins} = fIR_{Ins}(t) \quad \text{(Equation S74)}$$

where BW is body mass sent to the GLP-1RA module; while $f\_IR_{Uid}$, $fIR_{EGP}$, and $fIR_{Ins}$ are dimensionless metabolic-dysregulation functions supplied to the glucose–insulin module.

The two parameters fitted in this module are presented below:

| **Parameter** | **Fitted value best (min-max)** | **Unit** |
|---|---|---|
| λ_delayK | 0.44 (0.30 - 1.00) | 1/day |
| DiseaseProgression_Scaling | 0.35 (0.35 - 0.69) | fold-increase |

The remaining parameters were not fitted to data and were left unchanged:

| **Parameter** | **Used value (male)** | **Used value (female)** | **Unit** |
|---|---|---|---|
| cEI | 10.00 | | kcal/day |
| γ_FM | 0.03 | | kcal/(g·day) |
| γ_FFM | 0.15 | | kcal/(g·day) |
| π_FM | 0.18 | | kcal/g |
| π_FFM | 0.23 | | kcal/g |
| β | 0.40 | | dimensionless |

| | | | |
|---|---|---|---|
| K | 1.40 | | kcal/day |
| ρ_FM | 9.40 | | kcal/g |
| alpha_c | 0.10 | 0.13 | dimensionless |
| alpha_d | 0.00019 | 0.02 | dimensionless |
| alpha_k | 0.45 | 0.09 | 1/g |
| b_CLGI | 4.80 | | dimensionless |
| b_EGP | 70 | | dimensionless |
| b_Ins | 3.87 | | dimensionless |

### References


1. Guo J, Hall KD. Predicting Changes of Body Weight, Body Fat, Energy Expenditure and Metabolic Fuel Selection in C57BL/6 Mice. PLOS ONE. 2011 Jan 5;6(1):e15961. doi:10.1371/journal.pone.0015961

2. Gennemark P, Jansson-Löfmark R, Hyberg G, Wigstrand M, Kakol-Palm D, Håkansson P, et al. A modeling approach for compounds affecting body composition. J Pharmacokinet Pharmacodyn. 2013 Dec;40(6):651–67. doi:10.1007/s10928-013-9337-x PubMed PMID: 24158456.

3. Simonsson C, Lövfors W, Bergqvist N, Nyman E, Gennemark P, Stenkula KG, et al. A multi-scale *in silico* mouse model for diet-induced insulin resistance. Biochemical Engineering Journal. 2023 Feb 1;191:108798. doi:10.1016/j.bej.2022.108798

4. Silfvergren O, Rigal S, Schimek K, Simonsson C, Kanebratt KP, Forschler F, et al. M4 drug discovery: Human drug predictions from integrated preclinical insights exemplified with a GLP-1R agonist. European Journal of Pharmaceutical Sciences. 2026 Aug 30;107648. doi:10.1016/j.ejps.2026.107648

5. Nyman E, Brännmark C, Palmér R, Brugård J, Nyström FH, Strålfors P, et al. A hierarchical whole-body modeling approach elucidates the link between in Vitro insulin signaling and in Vivo glucose homeostasis. J Biol Chem. 2011 Jul 22;286(29):26028–41. doi:10.1074/jbc.M110.188987 PubMed PMID: 21572040; PubMed Central PMCID: PMC3138269.

6. Herrgårdh T, Li H, Nyman E, Cedersund G. An Updated Organ-Based Multi-Level Model for Glucose Homeostasis: Organ Distributions, Timing, and Impact of Blood Flow. Front Physiol. 2021 Jun 1;12. doi:10.3389/fphys.2021.619254

7. Silfvergren O, Simonsson C, Ekstedt M, Lundberg P, Gennemark P, Cedersund G. Digital twin predicting diet response before and after long-term fasting. PLOS Computational Biology. 2022 Sep 12;18(9):e1010469. doi:10.1371/journal.pcbi.1010469

8. Kleiber M. Body size and metabolic rate. Physiol Rev. 1947 Oct;27(4):511–41. doi:10.1152/physrev.1947.27.4.511 PubMed PMID: 20267758.
